# RNase III-mediated processing of a *trans*-acting bacterial sRNA and its *cis*-encoded antagonist

**DOI:** 10.1101/2021.03.19.434396

**Authors:** Sarah L. Svensson, Cynthia M. Sharma

**Affiliations:** Department of Molecular Infection Biology II, Institute of Molecular Infection Biology, University of Würzburg, 97080 Würzburg, Germany

**Keywords:** small regulatory RNA, sRNA, pathogenesis, post-transcriptional regulation, RNase III, *Campylobacter jejuni*

## Abstract

Bacterial small RNAs (sRNAs) are important post-transcriptional regulators in stress responses and virulence. They can be derived from an expanding list of genomic contexts, such as processing from parental transcripts by RNase E. The role of RNase III in sRNA biogenesis is less well understood despite its well-known roles in rRNA processing, RNA decay, and cleavage of sRNA-mRNA duplexes. Here, we show that RNase III processes a pair of *cis*-encoded sRNAs (CJnc190 and CJnc180) of the foodborne pathogen *Campylobacter jejuni*. While CJnc180 processing by RNase III requires CJnc190, In contrast, RNase III processes CJnc190 independent of CJnc180 via cleavage of an intramolecular duplex. We also show that CJnc190 directly represses translation of the colonization factor PtmG by targeting a G-rich ribosome binding site, and uncover that CJnc180 is a *cis*-acting antagonist of CJnc190, indirectly affecting *ptmG* regulation. Our study highlights a role for RNase III in sRNA biogenesis and adds *cis*-encoded RNAs to the expanding diversity of transcripts that antagonize bacterial sRNAs.

## INTRODUCTION

Bacterial small, regulatory RNAs (sRNAs) are an important class of post-transcriptional gene expression regulators that control adaptation to changing environmental conditions or stresses (Storz et al., 2011), or can also regulate virulence genes in pathogens (Quereda and Cossart, 2017; Svensson and Sharma, 2016; Westermann, 2018). They are also intimately associated with RNA-binding proteins (RBPs), such as RNA chaperones, as well as ribonucleases that are required for their maturation, stability, degradation, and/or function (Holmqvist and Vogel, 2018; Quendera et al., 2020). While most of the first identified and characterized sRNAs in bacterial genomes are expressed as stand-alone, intergenically-encoded transcripts, genome-wide RNA-seq based approaches have identified sRNAs hidden in unexpected genomic contexts, including 5′/3′ untranslated regions (UTRs) and coding regions of mRNAs, or even in housekeeping RNAs (reviewed in (Adams and Storz, 2020)). These include members of the expanding class of 3’ UTR-derived sRNAs mainly studied in Gammaproteobacteria (Miyakoshi et al., 2015a). These can be transcribed from an independent promoter, such as *E. coli* MicL (Chao et al., 2012; Guo et al., 2014), or can be processed from mRNAs by the single-stranded RNA endonuclease RNase E (Chao and Vogel, 2016; De Mets et al., 2019). Intergenically-encoded, stand-alone sRNAs can also require maturation by RNase E to increase their stability (Chae et al., 2011; Davis and Waldor, 2007; Hör et al., 2020), expose their seed regions (Papenfort et al., 2009), or even create two sRNAs with distinct regulons (Fröhlich et al., 2016; Papenfort et al., 2015).

Despite progress in defining mechanisms and regulatory consequences of complex sRNA biogenesis pathways in model Gammaproteobacteria, less is known about how such sRNAs are generated in bacteria lacking RNase E. RNase E has the most well-characterized role in bacteria (Bandyra and Luisi, 2018; Miyakoshi et al., 2015a) but is absent in ∼1/5 sequenced strains (Hui et al., 2014). In contrast, bacteria almost universally encode RNase III (Court et al., 2013). RNase III recognizes double-stranded RNA (11-20 base pairs long) in a mostly sequence-independent manner, and cleaves both strands to generate characteristic 2-3 nucleotide 3’-overhangs. Single-strand nicking, especially at imperfect duplexes, can also occur (Altuvia et al., 2018; Court et al., 2013; Le Rhun et al., 2017). Bacterial RNase III is mainly known for its role in rRNA processing, maturation or decay of certain mRNAs, and cleavage of sRNA-mRNA duplexes (Court et al., 2013). While the RNase III domain-containing Dicer and Drosha play a central role in sRNA biogenesis in eukaryotes (Carthew and Sontheimer, 2009), its role in sRNA biogenesis in bacteria is less clear. In *S. aureus*, RNase III generates RsaC sRNA from the 3′ UTR of the *mntABC* mRNA (Faubladier et al., 1990; Lalaouna et al., 2019). Moreover, its expropriation for biogenesis of CRISPR RNAs (Deltcheva et al., 2011; Dugar et al., 2018), as well as genome-wide studies of the RNase III targetome that report sRNAs as potential targets in Gram-negative as well as Gram-positive species (Altuvia et al., 2018; Gordon et al., 2017; Le Rhun et al., 2017; Lioliou et al., 2013, 2012; Lybecker et al., 2014; Rath et al., 2017), indicate that RNase III might process sRNAs in diverse bacteria. However, most of these remain to be validated or studied. In addition to an expanding genomic context of regulatory RNA sources, there is also emerging evidence of a high complexity of bacterial post-transcriptional networks involving not only cross-talk with transcriptional control, but also RNA antagonists that can sequester and modulate RNA-binding proteins (Dugar et al., 2016; Romeo and Babitzke, 2018; Sterzenbach et al., 2013; Wassarman, 2018) or even regulate stability or function of other RNAs as so-called competing endogenous RNAs (ceRNAs), RNA decoys/predators, or sponge RNAs (Figueroa-Bossi and Bossi, 2018; Grüll and Massé, 2019; Kavita et al., 2018). Such RNA antagonists can be derived from diverse cellular transcripts, including mRNAs (UTRs and coding regions) (Adams and Storz, 2020; Adams et al., 2021; Figueroa-Bossi et al., 2009; Miyakoshi et al., 2015b) and tRNAs (Lalaouna et al., 2015), or can be stand-alone sRNAs encoded in the core genome or in prophages (Bronesky et al., 2019; Melamed et al., 2020; Tree et al., 2014). Unbiased global biochemical and genetic screens for sRNA expression and regulation have recently recovered several characterized examples of *trans*-acting sRNA antagonists in Gram-positive and Gram-negative bacteria (Bronesky et al., 2019; Chen et al., 2021; Durand et al., 2021; Melamed et al., 2020), including those affecting infection phenotypes via antagonism of central sRNA regulators of virulence (Le Huyen et al., 2021). Despite reports of extensive antisense transcription in diverse bacteria and the demonstrated role of asRNAs in control of mRNA translation and stability (Thomason and Storz, 2010), less is known about whether RNAs encoded in *cis* to other sRNAs can act also as antagonists or how they might affect the biogenesis, stability, or function of their antisense sRNA partners.

The zoonotic food-borne pathogen *Campylobacter jejuni* is currently the leading cause of bacterial foodborne gastroenteritis worldwide (Burnham and Hendrixson, 2018; Havelaar et al., 2015). So far, how *C. jejuni* regulates its gene expression to adapt to different environments is unclear, as its genome encodes only three sigma factors (Parkhill et al., 2000; Young et al., 2007) and lacks homologs of certain global stress response regulators as well as of the global RNA chaperones Hfq and ProQ (Pernitzsch and Sharma, 2012; Quendera et al., 2020). Our comparative differential RNA-seq (dRNA-seq) analysis of multiple *C. jejuni* strains revealed many conserved and strain-specific sRNAs and asRNAs (Dugar et al., 2013). However, functions are largely unknown for most of these. Besides a so far missing general sRNA chaperone, it is also largely unknown which RNases participate in sRNA biogenesis and function in *C. jejuni*. Although *C. jejuni* is a Gram-negative, Epsilonproteobacteria surprisingly encode an RNase repertoire more similar to Gram-positives: for example, RNase Y and RNase J instead of RNase E (Parkhill et al., 2000; Pernitzsch and Sharma, 2012; Tomb et al., 1997). *C. jejuni* also encodes an RNase III homolog (Haddad et al., 2013), which participates in the biogenesis of Type II-C CRISPR RNAs (Dugar et al., 2018, 2013). Beyond this and a role in rRNA biogenesis, the function of *C. jejuni* RNase III is unclear.

Here we have characterized the biogenesis and mode of action of a conserved, processed pair of *C. jejuni cis*-encoded, antisense RNAs, CJnc180/190. We previously reported that deletion of these sRNAs affects *C. jejuni* virulence in a three-dimensional tissue-engineered model of the human intestine (Alzheimer et al., 2020). While this seemed to be at least in part mediated via repression of the flagellin modification factor PtmG, the underlying molecular mechanisms of this, the roles of the two sRNAs in PtmG regulation, as well as how they are processed remained unknown. Here, we demonstrate that both RNAs are processed by RNase III and that mature CJnc190 directly represses translation of *ptmG* mRNA by base-pairing with a G-rich sequence over its RBS. Surprisingly, although both RNAs are expressed antisense to each other, suggesting co-processing by RNase III, only CJnc180 requires its antisense partner for maturation. Instead, CJnc190 is transcribed as longer precursors which can fold into extended stem-loop structures that are processed independently of CJnc180 by RNase III. Finally, we demonstrate that CJnc180 is a *cis*-acting antagonist of CJnc190. Overall, our characterization of the CJnc190 biogenesis pathway demonstrates a role for RNase III in sRNA maturation and also reveals the potential for *cis*-encoded sRNA-sRNA targeting.

## RESULTS

### A product of the CJnc180/190 sRNA locus represses expression of *ptmG*

Our comparative dRNA-seq study of multiple *C. jejuni* isolates revealed a conserved pair of antisense sRNAs, CJnc180 and CJnc190 (annotated as 99 and 216 nucleotides (nt), respectively, in strain NCTC11168) (Dugar et al., 2013) (Figure 1A). RNA-seq patterns and northern blot analysis suggested both sRNAs might be processed from longer forms (Dugar et al., 2013). While CJnc180 was detected in wild-type (WT) as both a ∼90 nt “mature” 5′ end-derived species and a ∼160 nt longer putative precursor (pre-CJnc180) on northern blots, only a single CJnc190 RNA (∼70 nt) was detected. This CJnc190 species appeared to arise from the 3′ end of its annotated primary transcript (pre-CJnc190). Mature CJnc180 (hereafter, CJnc180) shows almost complete complementarity to mature CJnc190 (hereafter, CJnc190, Figure 1A).

**Figure 1.**
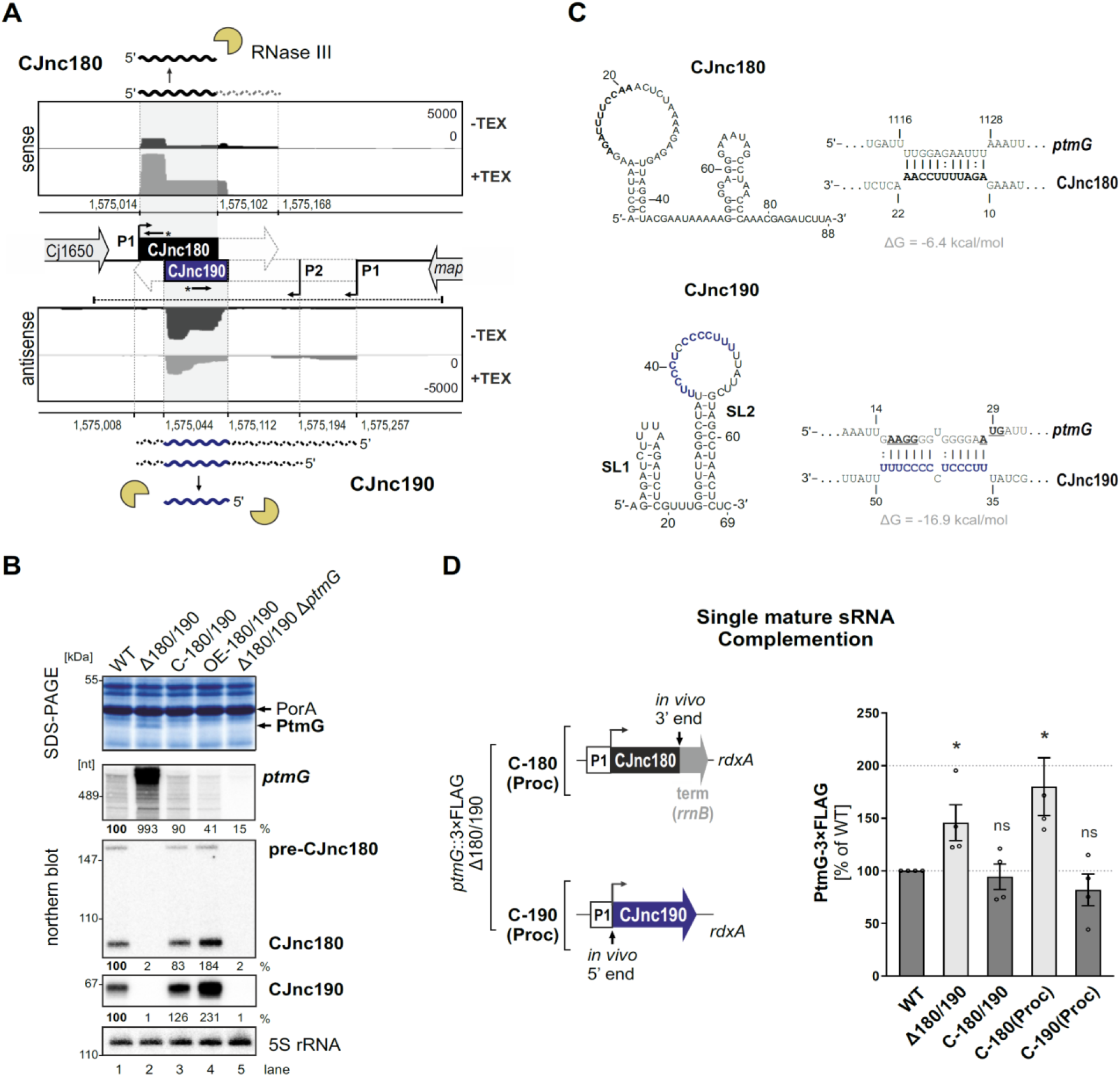
The processed CJnc190 sRNA represses *ptmG* expression. **(A)** dRNA-seq coverage (Dugar et al., 2013) for CJnc180 and CJnc190 sRNAs in *C. jejuni* NCTC11168. -/+TEX: mock/treated terminator exonuclease (TEX) dRNA-seq libraries. Treatment degrades processed (non-triphosphorylated) 5’ ends, enriching 5’-triphosphorylated primary transcript ends at transcription start sites (TSS). Bent arrows: TSS. Black dashed line: genomic region used for complementation (C-180/190). Starred arrows: northern blot probes for mature sRNAs. P1/P2: promoter motifs. **(B)** SDS-PAGE and northern blot analyses of total protein and RNA, respectively, from *C. jejuni* WT and sRNA/*ptmG* mutant strains. Upregulated ∼45 kDa PtmG and non-regulated PorA control are indicated. Probes for the mature sRNAs (CSO-0189/0185 for CJnc180/190, respectively (starred arrows, panel A), and the 5’ end of the *ptmG* ORF (CSO-1666) were used. As a loading control, 5S rRNA was probed with CSO-0192. OE, second-copy overexpression. **(C)** Predicted secondary structures (RNAfold) (Lorenz et al., 2011) and *ptmG* interactions (IntaRNA) (Mann et al., 2017) for mature CJnc180 and CJnc190. Bold/blue: potential *ptmG* pairing residues for CJnc180/CJnc190, respectively. Underlined: *ptmG* RBS/start codon. **(D)** Complementation of *ptmG* regulation in Δ180/190 with single mature sRNAs. *(Left)* To express CJnc180 only (C-180(Proc)), its mature 3’ end was fused to the *E. coli rrnB* terminator; transcription is driven from its native promoter. For C-190(Proc), its mature 5′ end was fused to its annotated TSS and 125 upstream nucleotides (P1 promoter). (*Right)* PtmG-3×FLAG levels measured by western blotting. Error bars represent standard error of the mean (SEM) of four independent replicates. **\***: p<0.05, ns: not significant, vs. WT. See also **Supplementary Figure S4A**.

We previously observed that deletion of CJnc180/190 affects *C. jejuni* adherence and internalization in our Caco-2 cell based tissue-engineered model of the human intestine (Alzheimer et al. 2020), suggesting that CJnc180 and/or CJnc190 regulate genes involved in *C. jejuni* virulence. Analysis of total protein profiles by SDS-PAGE revealed an ∼45 kDa band upregulated upon deletion of CJnc180/190 (Δ180/190) (Figure 1B), which was identified as the colonization/infection-relevant flagellin modification factor PtmG (Cj1324) (Alzheimer et al., 2020; Howard et al., 2009) based on mass spectrometry (**Supplementary Figure S1; Supplementary File 1 - Table S1**). The upregulated band was no longer observed in a Δ180/190 Δ*ptmG* double mutant, confirming it as PtmG (Figure 1B). Complementation of Δ180/190 with a region spanning the Cj1650 and *map* intergenic region (C-180/190, dashed line, Figure 1A) at the unrelated *rdxA* locus rescued expression of both sRNAs and restored PtmG repression (Figure 1B).

Northern blot analysis further demonstrated that not only the protein level but also *ptmG* mRNA levels are upregulated almost 10-fold in Δ180/190 vs. WT and are reduced (2-fold) upon overexpression of CJnc180/190. Overall, these data indicate that at least one of the two sRNAs is involved in repression of the gene encoding the *ptmG* colonization factor (Alzheimer et al., 2020; Howard et al., 2009).

### The mature CJnc190 sRNA is sufficient to repress *ptmG*

To disentangle the roles of each sRNA in potential direct regulation of *ptmG*, we first defined the 5’ and 3’ ends of each mature sRNA in WT using primer extension and 3′RACE (rapid amplification of cDNA ends) (for details, see **Supplementary Figure S2 & S3)**. Subsequent predictions for secondary structures and potential *ptmG* mRNA interactions of the mature sRNAs revealed a strong potential for a C/U-rich loop within CJnc190 to base-pair with the ribosome binding site (RBS) and start codon of *ptmG* mRNA (Figure 1C). A less stable interaction (ΔG = −6.4 vs. −16.9 kcal/mol for CJnc190:*ptmG*) was predicted between CJnc180 and the 3′ end of the *ptmG* coding region, suggesting that CJnc190, and not CJnc180, directly represses *ptmG* translation. To further test this, we constructed Δ180/190 complementation strains expressing either mature CJnc180 or CJnc190 alone (hereafter C-180(Proc) or C-190(Proc)) from their annotated native promoters (P1, Figure 1D, *left*) and measured rescue of *ptmG* repression. A chromosomally epitope-tagged PtmG-3×FLAG fusion was upregulated 1.5-fold upon deletion of CJnc180/190 and rescued to WT levels in the C-180/190 complemented strain (Figure 1D, *right*). In line with the prediction that *ptmG* translation is repressed by base-pairing with CJnc190 and not CJnc180, the C-190(Proc) complementation strain had PtmG-3×FLAG levels comparable to WT, whereas C-180(Proc) did not restore regulation of PtmG-3×FLAG. A similar trend was seen for *ptmG* mRNA levels (**Supplementary Figure S4A**). Moreover, a translational GFP reporter fusion of the 5’UTR and first 10 codons of *ptmG* (*ptmG(10th)*-GFP) was repressed by CJnc190 when transcribed from either the native *ptmG* promoter or the unrelated σ^28^ (FliA)-dependent *flaA* promoter, confirming regulation at the post-transcriptional level (**Supplementary Figure S4B**). Collectively, these observations showed that CJnc180 is dispensable and mature CJnc190 is sufficient for post-transcriptional repression of *ptmG*.

### CJnc190 represses *ptmG* translation by base-pairing with its G-rich RBS

We next validated direct interaction between CJnc190 and *ptmG* mRNA and its requirement for regulation using *in vitro* and *in vivo* approaches. These experiments included compensatory base-pair exchanges within the predicted CJnc190-*ptmG* duplex (Figure 2A). Gel mobility shift assays using *in vitro*-transcribed RNAs showed that processed CJnc190 binds the *ptmG* leader, and that a single C-to-G change in its C/U-rich loop (M1) is sufficient to almost completely abolish complex formation (Figure 2B). Similarly, a single point mutation (M1′) in the *ptmG* 5′UTR also disrupted interaction with CJnc190, while introduction of the compensatory base exchange in CJnc190 (M1) restored binding. Mature CJnc180 did not bind *ptmG*, in line with the weak predicted CJnc180:*ptmG* interaction. Reciprocal experiments with labeled CJnc190 (WT/M1) and unlabeled *ptmG* leader (WT/M1′) further confirmed the interaction (**Supplementary Figure S5A**). Adding increasing amounts of unlabeled *ptmG* leader (WT) to labeled mature CJnc190 in Inline probing assays protected nucleotides in the C/U rich loop region of 5′ end labeled mature CJnc190 from cleavage, in agreement with the predicted interaction with the *ptmG* leader (Figures 2A **&** 2C). The same molar ratio of *ptmG* M1′ mutant leader showed less protection, indicating destabilization of the interaction. Reciprocal experiments with labeled WT *ptmG* leader and WT or M1 CJnc190 further confirmed the predicted interaction site on *ptmG* (**Supplementary Figure S5B**).

**Figure 2.**
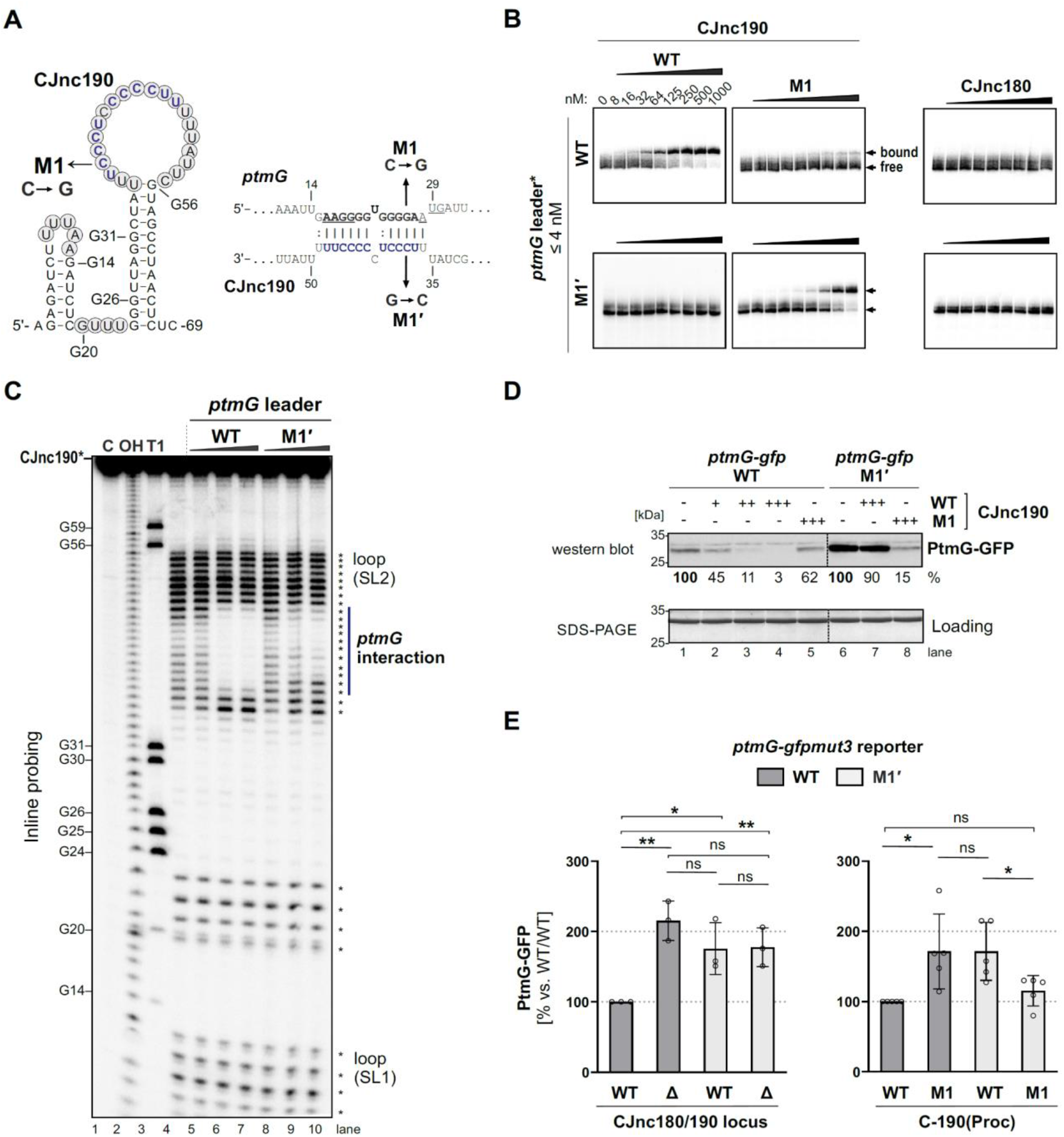
CJnc190 represses translation of *ptmG* via base-pairing with its G-rich RBS. **(A)** CJnc190 secondary structure based on Inline probing (C) and interaction with the *ptmG* leader showing mutations (M1/M1′) introduced into the interaction site. Circled residues: single-stranded regions mapped by Inline probing. Blue/bold residues: *ptmG*/CJnc190 nucleotides protected in Inline probing (panel A and **Supplementary Figure S5B**). RBS/start codon are underlined. **(B)** *In vitro* gel shift assay of ^32^P-5′-labeled (marked with *) *ptmG* leader (WT/M1′) with unlabeled CJnc190 WT/M1 as well as CJnc180 sRNAs. **(C)** Inline probing of 0.2 pmol ^32^P-5′-end-labeled CJnc190 sRNA in the absence or presence of 0.2/2/20 pmol unlabeled *ptmG* leader (WT/M1′). C - untreated control; T1 ladder - G residues (indicated on left); OH - all positions (alkaline hydrolysis). **(D)** *In vitro* translation of a *ptmG(10th)*-GFP reporter (5′ UTR and first 10 codons of *ptmG* fused to *gfpmut3*, 2 pmol) in an *E. coli* cell-free system +/- CJnc190 (WT/M1 +: 2 pmol, ++: 20 pmol, +++: 100 pmol) detected by western blotting with an anti-GFP antibody. A Coomassie-stained gel of the same samples served as a loading control. **(E)** PtmG(10th)-GFP (WT/M1′) reporter expression *in vivo* +/- mature CJnc190 (WT/M1) measured by western blot analysis. PtmG(10th)-GFP levels are the mean of three (left) or five (right) independent replicates, with error bars representing the SD. **: p<0.01, *: p<0.05, ns: not significant, vs. WT. See also **Supplementary Figure S5C**.

The observed base-pairing with a G-rich sequence at the *ptmG* RBS suggested that CJnc190 acts by repressing *ptmG* translation. Indeed, the addition of increasing molar ratios (1-, 10-, 50-fold; + to +++) of mature CJnc190 to a *ptmG(10th)-gfp* translational reporter mRNA in an *in vitro* translation system repressed GFP levels in a dose-dependent manner (Figure 2D). Consistent with the disrupted interaction, the M1 mutation in the CJnc190 loop partially restored translation of the reporter. While the *ptmG* M1′ mutation increased its translation compared to WT, already independent of CJnc190 addition, Inline probing experiments did not reveal marked differences in secondary structure for the native (non-GFP-fusion) WT and M1′ *ptmG* leaders *in vitro* (**Supplementary Figure S5B**). Nonetheless, while the addition of CJnc190 WT did not strongly affect translation of the mutant reporter, addition of CJnc190 M1, carrying the compensatory exchange in its C/U-rich loop, strongly reduced GFP levels generated from the M1′ reporter, indicating restored regulation (Figure 2D).

In line with the *in vitro* results, introduction of the M1′ mutation *in vivo* in the *ptmG(10th)*-GFP reporter fusion derepressed GFP levels, although not to those of an isogenic Δ180/190 strain (Figure 2E, *left*). To account for different levels of CJnc190 in the WT and C-190(Proc) strains, we next compared regulation of the reporter by WT/M1 CJnc190 in the C-190(Proc) background. In line with a disrupted interaction, CJnc190 M1 did not repress the WT *ptmG* reporter to the same levels as CJnc190 WT (Figure 2E, *right*), even though WT/M1 sRNAs were similarly expressed (**Supplementary Figure S5C**). Likewise, when WT CJnc190 was expressed with the *ptmG* M1′ reporter, GFP levels were also higher compared to the WT sRNA/leader strain. Finally, in the strain with the compensatory mutations combined (M1/M1′), GFP levels were similar to the isogenic WT/WT (Figure 2E). Together, our *in vitro* and *in vivo* experiments demonstrate that CJnc190 represses *ptmG* translation via base-pairing with its RBS.

### RNase III processes the CJnc180/190 sRNAs

Next, we set out to gain insight into the biogenesis of the CJnc180/190 sRNA pair. While deletions of most non-essential RNases/RNA degradation enzymes had no major impact on processing of the two RNAs (**Supplementary Figure S6A**), deletion of RNase III (Δ*rnc*) had a dramatic effect on both CJnc180 and CJnc190, abolishing accumulation of the mature sRNA species (Figure 3A). For CJnc180, the longer transcript of WT (pre-CJnc180, ∼160 nt) was still detected in Δ*rnc*, but at higher levels. For CJnc190, the mature form of WT was completely absent in Δ*rnc* and instead five longer “pre-CJnc190” species (∼150-280 nt) were detected. All CJnc180 and CJnc190 unprocessed species were absent in a Δ180/190 Δ*rnc* double mutant, ruling out cross-hybridization.

**Figure 3.**
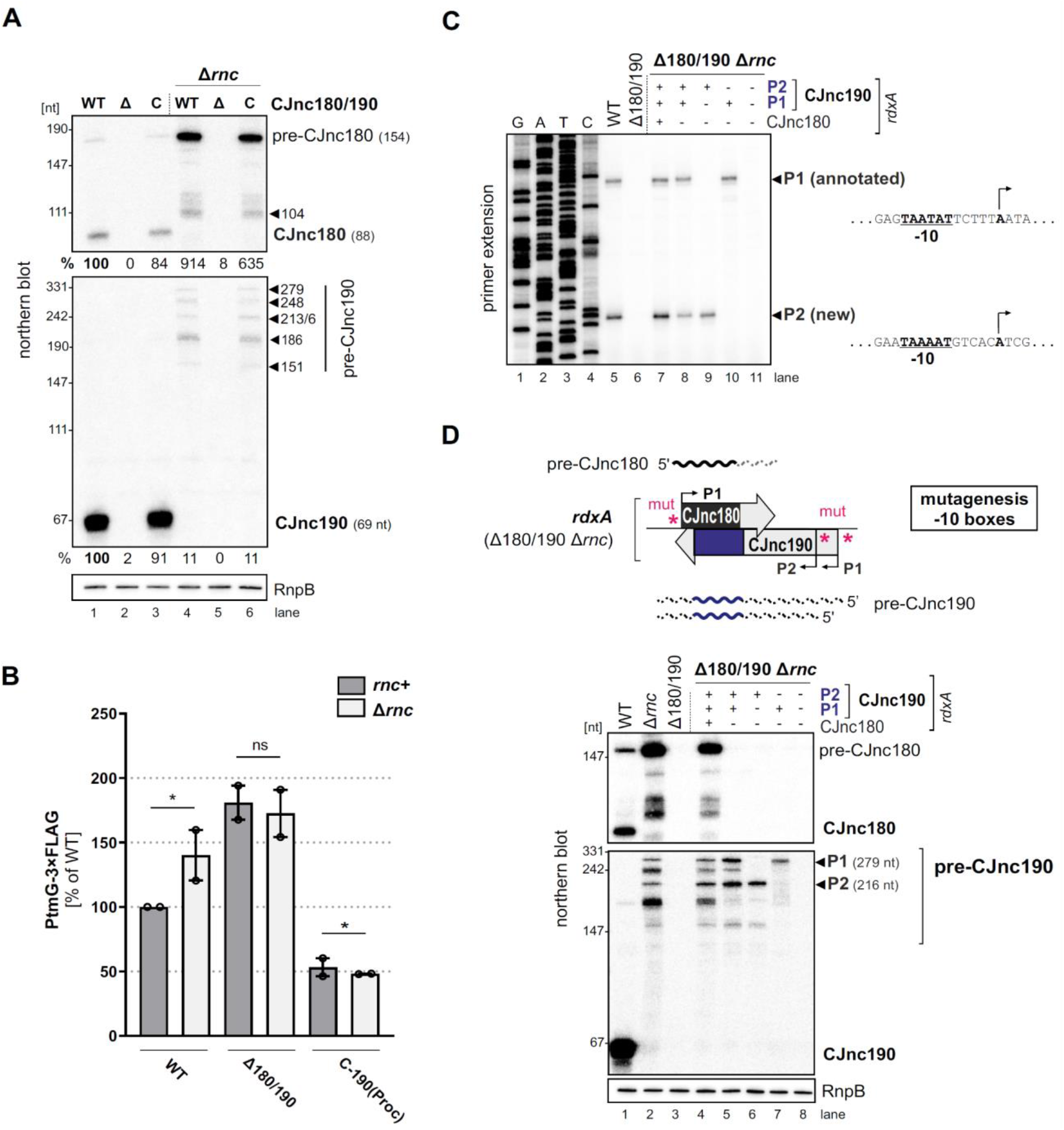
RNase III processes CJnc190 precursors expressed from two promoters. **(A)** Northern blot of CJnc180 and CJnc190 processing by RNase III in total RNA. Indicated lengths are based on primer extension and 3’ RACE. Quantification is for all bands combined. **(B)** Effect of *rnc* (RNase III) deletion on PtmG-3×FLAG levels in the absence or presence of CJnc180/190 sRNAs. Error bars represent the SEM of two independent replicates. *: p<0.1, ns: not significant, Student’s unpaired *t*-test. See also **Supplementary Figure S6B**. **(C)** Primer extension analysis of pre-CJnc190 5′ ends in WT and promoter mutant strains (Δ*rnc* background). Total RNA was annealed with the same probe for mature CJnc190 used for northern blots (CSO-0185). A sequencing ladder was generated with the same probe (lanes 1-4). P1/P2: putative CJnc190 primary transcripts/5’ ends. The full gel is shown in **Supplementary Figure S8. (D)** Validation of CJnc180/190 promoters. (*Top*) Strategy for testing CJnc180/190 promoter activity by complementation of Δ180/190 with -10 box mutant alleles at *rdxA.* “mut” - several point mutations introduced into the predicted -10 box (see **Supplementary File 1 - Table S5** for details). (*Bottom*) Northern blot analysis of pre-CJnc180/CJnc190 in sRNA promoter mutant strains (Δ*rnc* background). (-/+): promoter mutant/WT. Probes for the mature sRNAs were used (CSO-0189/0185, respectively, for CJnc180/190; Figure 1A). RnpB served as loading control (probed with CSO-0497). For primer extension analysis of the same strains, see panel C.

We next asked if RNase III processing of CJnc190 affects *ptmG* regulation. Deletion of *rnc* increased PtmG-3×FLAG protein and mRNA levels to those similar to Δ180/190 (Figure 3B & **Supplementary Figure S6B**), and CJnc180/190 deletion in Δ*rnc* did not increase levels further. In contrast to “native” CJnc190 (*i.e*., processed from precursors), the “mature” sRNA of the C-190(Proc) strain (transcribed directly from its mature 5′ end) was not markedly affected by *rnc* deletion (**Supplementary Figure S2B**) and was still capable of *ptmG* regulation in the *rnc* deletion background (Figure 3B). The mild increase in *ptmG* mRNA levels upon *rnc* deletion in C-190(Proc) suggests that RNase III could also play a minor role in cleavage of CJnc190:*ptmG* duplexes (**Supplementary Figure S6B**). The combined levels of all CJnc190 species were 10-fold lower in Δ*rnc* compared to WT (Figure 3A, compare lanes 1 and 4). Rifampicin stability assays showed that the CJnc180 precursor was stabilized in the Δ*rnc* mutant when compared to WT (**Supplementary Figure S6C**), which further validates processing by RNase III. Moreover, while the half-life of mature CJnc190 in the WT strain was >64 min, CJnc190 precursors had half-lives of 2-4 min in the Δ*rnc* mutant. Taken together, these results indicate that RNase III processes both sRNAs and affects *ptmG* regulation, likely by generating a more stable CJnc190 sRNA species.

### CJnc190 precursors are transcribed from two promoters

To understand the unique RNase III-mediated processing of CJnc180 and CJnc190, we next characterized their precursors. Further northern blot analysis of total RNA from Rnc+ and Δ*rnc* strains with diverse probes showed that the 3′ end of CJnc180 is removed by RNase III processing, and that its 5’ end is RNase III-independent (**Supplementary Figure S7A & 7B**). Additional probing for CJnc190 in WT and Δ*rnc* suggested that they differ in both their 5’ and 3’ ends (**Supplementary Figure S7A & 7C**). We used primer extension to map the 5’ ends of the Δ*rnc* precursors for both sRNAs. While CJnc180 had a single, RNase III-independent 5’ end that mapped to its annotated TSS (**Supplementary Figure S2A**), we detected two CJnc190 5’ ends in a Δ*rnc* background (Figure 3C, lane 5). One end matched the annotated primary TSS for CJnc190 (Dugar et al., 2013), which has a putative -10 box (TAATAT) immediately upstream (Figure 3C, right). Inspection of the nucleotides upstream of the second detected 5’ end also revealed a near consensus -10 box (TAAAAT). This suggested that CJnc190 precursors might be transcribed from at least two sigma70-dependent promoters.

To validate the activity of the three putative CJnc180/190 promoters [annotated CJnc180(P1) and CJnc190(P1); predicted CJnc190(P2)] *in vivo*, we performed site-directed mutagenesis of their respective -10 boxes in the C-180/190 complementation construct (Figure 3D, for details see methods and **Supplementary File 1 - Table S5**). Analysis of CJnc180 in a Δ*rnc* background in the generated promoter mutant strains showed that disruption of the annotated CJnc180 promoter -10 box (strain C-190 only) fully abolished CJnc180 expression (Figure 3D, lane 5), confirming its single TSS. CJnc180 promoter disruption reduced accumulation of some CJnc190 precursors (but did not affect their 5′ ends), suggesting transcriptional interference might impact 3′ ends. Next, we inspected pre-CJnc190 expression in strains with disruptions in either CJnc190(P1) or CJnc190(P2) added to the C-190 only construct. In a strain with CJnc180 and CJnc190 P1 promoters inactivated, the two longest CJnc190 species of Δ*rnc* were absent, and detection of a shorter CJnc190 transcript supported the presence of a second promoter (Figure 3D, lane 6). In the strain with only CJnc190 P1 active, instead only a longer (∼280 nt) transcript was detected (lane 7). Finally, disruption of all three putative promoters (C-3×mut) completely abolished expression of all CJnc180 and CJnc190 transcripts (lane 8). Primer extension analysis of RNA from the same promoter mutant strains validated that the two 5’ ends we detected were dependent on the upstream -10 sequences (Figure 3C, lanes 7-11). Overall, mapping of CJnc180 and CJnc190 precursors in Δ*rnc* suggested that mature CJnc180 is transcribed from a single promoter and derived from its precursor 5′ end, while mature CJnc190 is generated from the middle of transcripts with different 3′ end lengths arising from two promoters.

### CJnc180, but not CJnc190, requires its antisense partner for processing by RNase III

As the RNase III-dependent biosynthesis pathway of CJnc190 and CJnc180 makes them distinct from other processed sRNAs characterized in Gammaproteobacteria, we next explored their maturation in more detail. Because of their extensive complementarity, we hypothesized that RNase III coprocesses CJnc180:CJnc190 duplexes. To examine this, we repeated northern blot analysis of the CJnc180/190 promoter-inactivation allele strains, but this time in an RNase III+ background. This surprisingly showed that the C-190 only strain (with the CJnc180 promoter disrupted) still expressed mature CJnc190 (Figure 4A). We also found that CJnc190 precursors from either P1 or P2 could give rise to mature CJnc190, although when expressed from P1 or P2 alone, mature sRNA levels were approximately 60% of those detected in the WT, C-180/190, or C-190 only strains. This suggests that both promoters drive transcription of pre-CJnc190 precursors and their combined activity in exponential phase contributes to levels of the mature sRNA.

**Figure 4.**
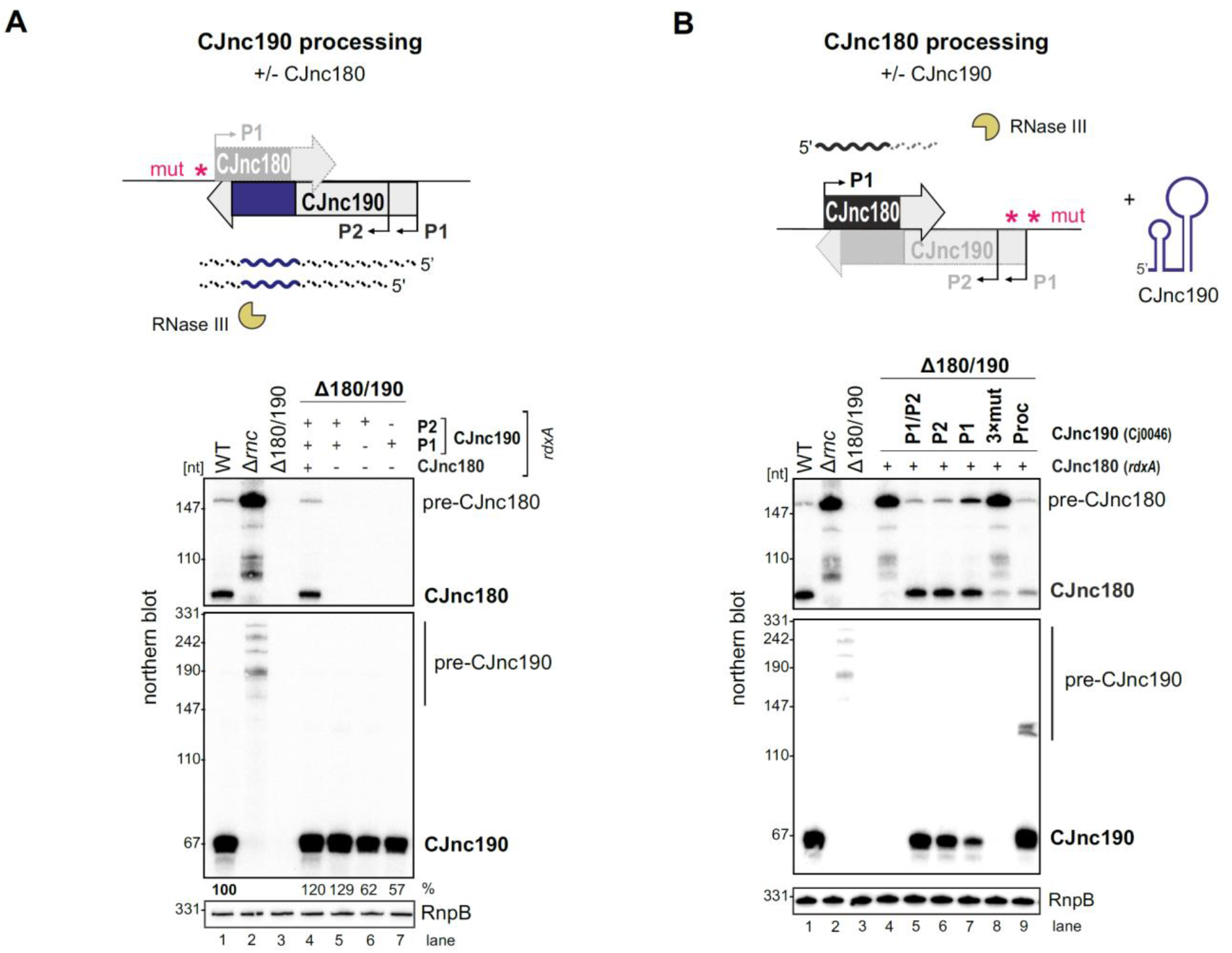
CJnc180 requires its antisense partner for RNase III-mediated processing, while CJnc190 processing is CJnc180-independent. Top of both panels: Approach for testing processing in the absence of the antisense partner using promoter mutant alleles. “mut” - several point mutations introduced into the -10 box (see **Supplementary File 1 - Table S5** for details). **(A)** Northern blot analysis of pre-CJnc190 processing *in vivo* +/- CJnc180. The Δ180/190 strain was complemented at *rdxA* with WT or CJnc180/190 promoter mutant alleles in an *rnc*+ background. (+/-) indicates if a promoter in the CJnc180/190 allele is WT/mutant. **(B)** Pre-CJnc180 processing in the presence or absence of CJnc190. Pre-CJnc180 was introduced into *rdxA* of a Δ180/190 strain. Different CJnc190 species were expressed from the unrelated Cj0046 pseudogene locus. For northern blot detection of CJnc180 and CJnc190, probes for the mature sRNAs were used (CSO-0189 and CSO-0185, respectively). RnpB (probed with CSO-0497) served as a loading control.

We next examined whether CJnc180 processing by RNase III is likewise CJnc190-independent. Complementation of Δ180/190 with pre-CJnc180 alone (strain C-180; both CJnc190 promoters disrupted) surprisingly revealed that in contrast to CJnc190, CJnc180 was not processed without its antisense partner (Figure 4B, lane 4). Instead, as in Δ*rnc,* we detected only pre-CJnc180. To confirm that CJnc180 processing requires pairing with CJnc190, we added back different CJnc190 species to the second unrelated Cj0046 pseudogene locus in the C-180 only strain (lanes 5-7). Expression of CJnc190 (either P1 or P2) from this second locus restored processing of CJnc180, and even in *trans* expression of “mature” CJnc190 from the C-190(Proc) strain (which overlaps the CJnc180 3′ end) was sufficient to restore CJnc180 processing (lane 9). Overall, these mutational analyses indicate that while CJnc190 is processed independently of CJnc180, CJnc180 processing is mediated by expression of CJnc190 in *cis* or *trans*.

### CJnc190 processing is mediated by an intramolecular duplex

We hypothesized that CJnc190 processing by RNase III might be mediated by A) a second *trans*-encoded RNA, or B) via cleavage of an intramolecular duplex. Consistent with the second hypothesis, we detected diverse 3’ ends for CJnc190 by northern blotting (**Supplementary Figure S7**). We next mapped the 3’ ends of CJnc190 precursors in Δ*rnc* by 3’RACE. The combined northern blot, primer extension, and 3’RACE information suggested that the most abundant CJnc190 precursor species (∼186 nt) arises from P2 with a 3′ extension beyond mature CJnc190 (**Supplementary Figure S3 & S7**). We performed folding predictions for six precursors supported by our 5′ and 3′ mapping (**Supplementary Figure S9**). The most abundant CJnc190 precursor (pre-CJnc190, 186 nt from P2) can fold into a long duplex flanking the mature sRNA, with an additional 5′ hairpin (Figure 5A, *left*). While we also detected RNase III cleavage sites within the 5’ hairpin by primer extension (A/A’ in Figure 5A; **Supplementary Figure S9**), deletion of this region did not affect maturation of CJnc190 (**Supplementary Figure S10**). Moreover, these positions do not reflect the mature CJnc190 5′ end. This indicates that the 5’ hairpin of pre-CJnc190 species is not required for processing.

**Figure 5.**
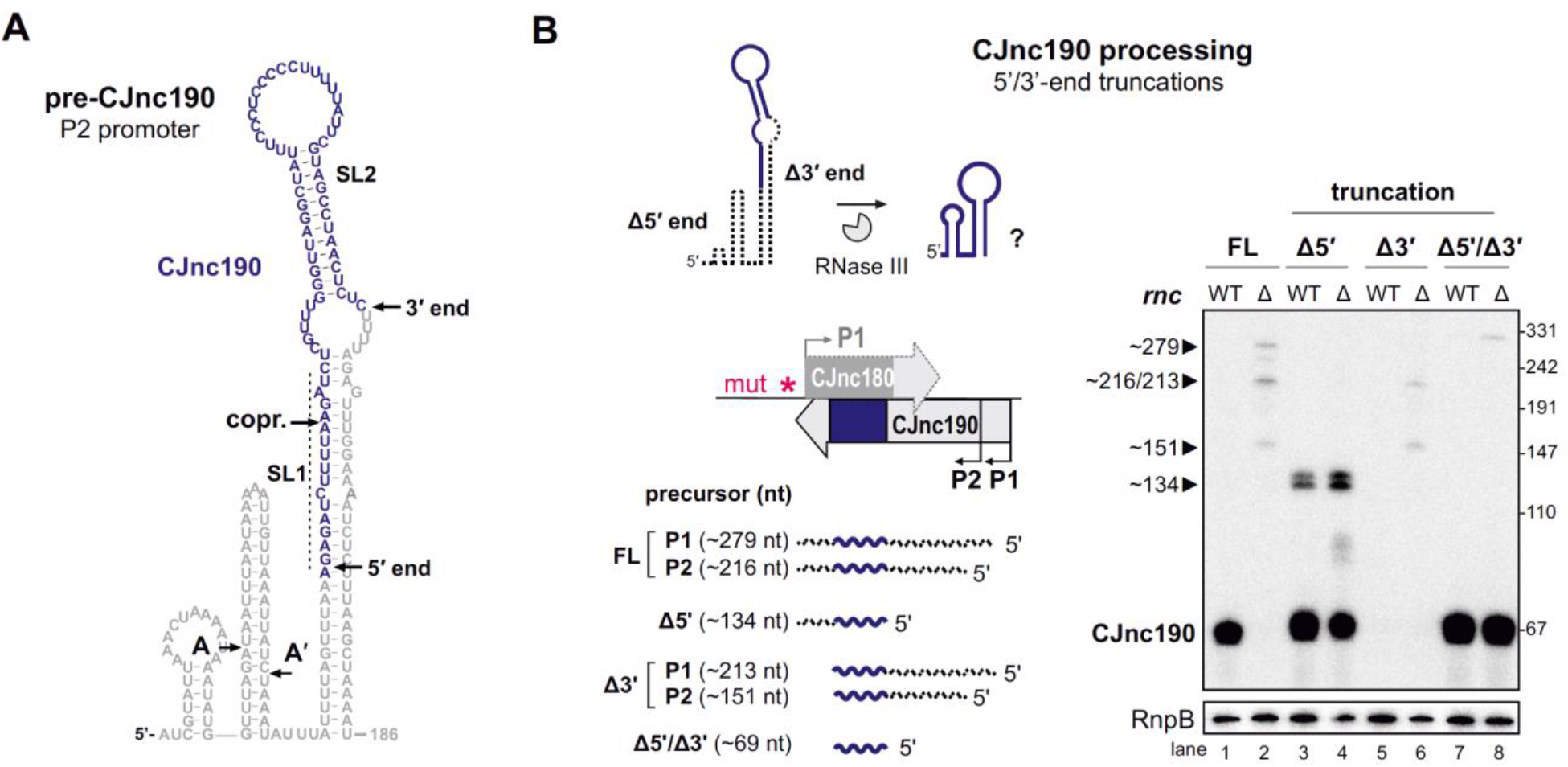
Intramolecular duplex-mediated processing of CJnc190. **(A)** Predicted secondary structure of the 186 nt pre-CJnc190 precursor transcribed from P2. Blue residues: mature sRNA. A/A’: putative intermediate 5’ ends identified by primer extension. Residues of the mature sRNA are blue. A putative co-processing site (copr.) with CJnc180 (see Figure 6A, *bottom*) is indicated. **(B)** Both 5′ and 3′ ends of CJnc190 are required for processing *in vivo*. *Left*: CJnc190 (without CJnc180) was expressed from the *rdxA* locus in Δ180/190 as full-length (FL) or as three versions with truncations (Δ) at the mature sRNA ends (5’ end, 3’ end, or both). For 5’ truncations, CJnc190 was fused to its P1 promoter. “mut” - several point mutations introduced into the predicted -10 box (see **Supplementary File 1 - Table S5** for details). (*Right*) CJnc190 expression and processing was detected by northern blotting with a probe for the mature sRNA (CSO-0185), while RnpB (CSO-0497) served as a loading control. Total RNA from strains expressing CJnc190 versions on the left were analyzed in an Rnc+ (WT) or Δ*rnc* background. The expected size of each unprocessed precursor is indicated on the left.

We therefore next examined the requirement of the extended 5′-3′ end duplex flanking the mature sRNA for processing. To test if the 3’ end is required for processing by RNase III, we truncated the 3′ end of pre-CJnc190 to the position of the mature sRNA (Δ3’). Unlike a 5′ end truncation to the mature CJnc190 end, this 3’ end truncation abolished detection of mature CJnc190 (Figure 5B, lanes 3/4 & 5/6). Removal of the same 3’ region also abolished truncation of either of the 5’-hairpin-truncated species (A or A’) (**Supplementary Figure S10B,** lanes 5/6 & 9/10). In contrast, the mature sRNA was detected when the 3′ end was removed from the 5′-truncated version (Figure 5B, lanes 7 & 8), suggesting the unprocessed sRNA is not stable. Only CJnc190 precursors with a 3′ extension beyond the mature sRNA showed base-pairing between 5′ and 3′ precursor ends (**Supplementary Figure S9**). Together, this suggests that RNase III cleaves at both ends of the precursor to generate mature CJnc190, and provides insight into how its processing is independent of CJnc180. However, based on mature CJnc190 ends detected in WT, the processing steps following RNase III cleavage remain to be determined.

### CJnc190 drives processing of CJnc180

We next examined why CJnc180 was not processed without CJnc190. We mapped the 3’ ends of pre-CJnc180 to positions corresponding to ∼104 and ∼154 nts (major product), consistent with northern blots (Figure 3A & **Supplementary Figure S3**). Compared to pre-CJnc190, the predicted secondary structure of pre-CJnc180 species (Figure 6A) or 104 nt pre-CJnc180 (**Supplementary Figure S11A**) do not contain distinct long duplexes like pre-CJnc190. We next performed *in vitro* RNase III cleavage assays with T7-transcribed pre-CJnc180 (154 nt) in the absence or presence of mature CJnc190 to see if we could recapitulate processing *in vitro*. In contrast to *in vivo*, RNase III could cleave pre-CJnc180 even without CJnc190, although the cleavage site was located ∼100 nt from the 5′ end (Figure 6B, asterisk), rather than at the RNase III-dependent 3′ end of the mature sRNA detected *in vivo* (Figure 6A). In contrast, reactions with increasing amounts of mature CJnc190 generated the *in vivo* RNase III-dependent cleavage site/3′ end (Figure 6B), as well as a second site (**Supplementary Figure S11**, double asterisk). Together, our data suggest that while RNase III is sufficient for maturation of structured CJnc190, CJnc180 also requires its antisense partner for processing.

**Figure 6.**
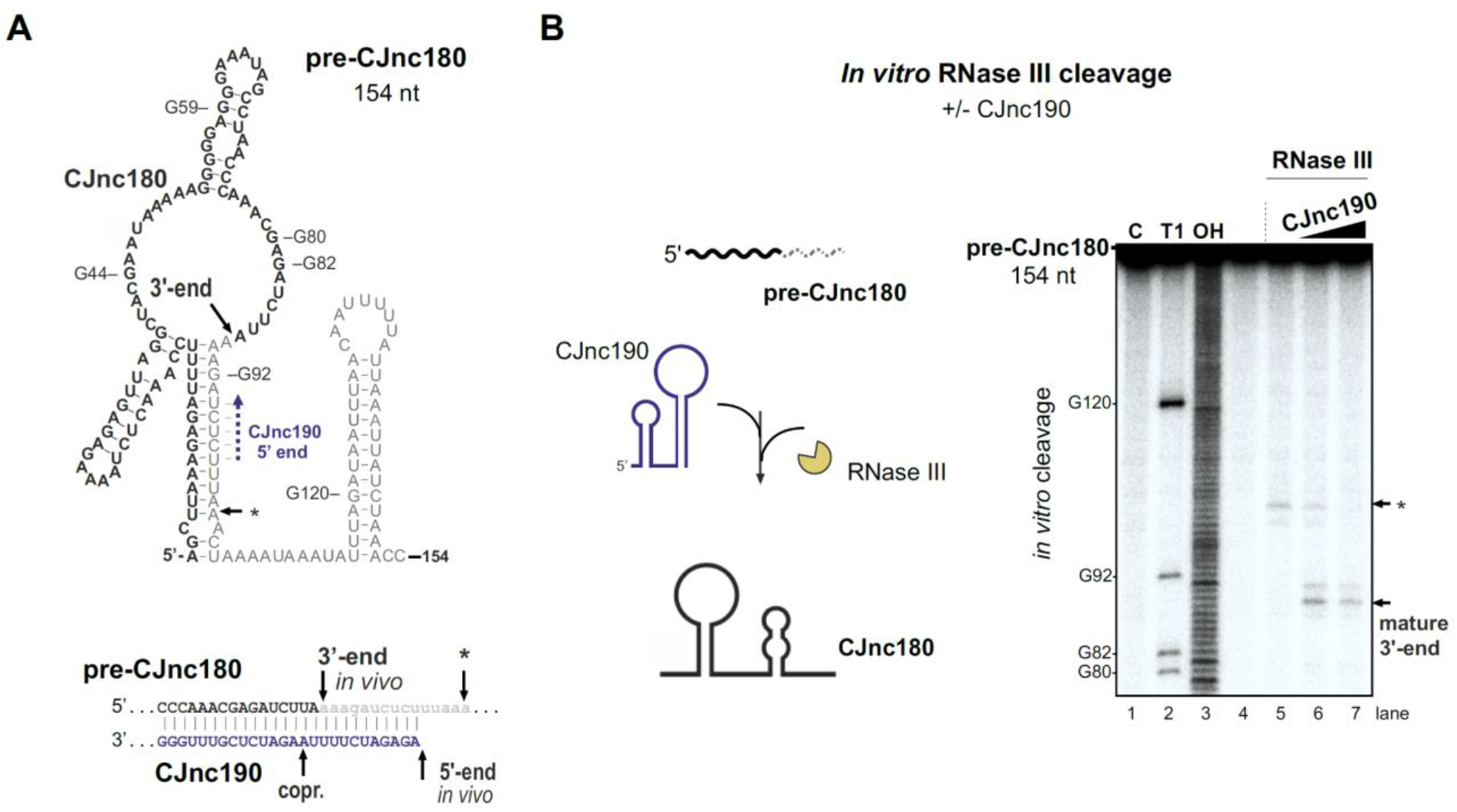
CJnc190 drives processing of CJnc180 by RNase III. **(A)** Predicted secondary structure of the pre-CJnc180 precursor (154 nt). Residues of the mature sRNA are black. The CJnc190-dependent CJnc180 3′ end is indicated. Asterisk: CJnc190-independent *in vitro* cleavage site (panel B). Blue dashed arrow: Region of potential base-pairing with 5′ end of mature CJnc190. (*Bottom*) Potential co-processing of the predicted CJnc190 (mature) and pre-CJnc180 duplex. **(B)** *In vitro* RNase III cleavage of ^32^P-labeled (5’ end) pre-CJnc180 (154 nt). *(Left)* ^32^P-5’-end labeled *in vitro* transcript (0.2 pmol) was incubated in the presence or absence of unlabeled mature CJnc190 (0.2 or 2 pmol) and subjected to cleavage with RNase III. (*Right*) Cleavage products were separated on a denaturing gel. C - untreated control; T1 ladder - G residues (indicated on left); OH - all positions (alkaline hydrolysis). The full gel is shown in **Supplementary Figure S11B**.

### CJnc180 indirectly affects *ptmG* via CJnc190 antagonism

While we found that CJnc180 was not required for CJnc190 processing, several *C. jejuni* and *C. coli* strains express this antisense RNA (Dugar et al., 2013; Riedel et al., 2020), suggesting it has a conserved function. Because of its extensive complementarity to CJnc190, as well as its co-processing, we hypothesized that it might serve as a CJnc190 antagonist and indirectly affect *ptmG* regulation. We therefore examined the effect of CJnc180 overexpression on PtmG regulation. Overexpression of full-length CJnc180 or “pre-processed” CJnc180(Proc) increased PtmG-3×FLAG to levels intermediate between Δ180/190 and WT (Figure 7A). In WT, the ratio of CJnc190 to all CJnc180 transcripts in log phase was ∼35:1 (**Supplementary Figure S12B**). CJnc180 overexpression decreased this ratio to ∼13:1. This suggested that the CJnc180 antisense RNA can in fact influence *ptmG* indirectly via an effect on CJnc190. In contrast to its overexpression, abolishing CJnc180 expression in log phase did not significantly affect PtmG-3×FLAG levels when CJnc190 was expressed from both P1 and P2 (Figure 7B). However, when we expressed CJnc190 from only a single promoter (P1 or P2), thereby reducing its overall levels, the absence of CJnc180 expression significantly affected *ptmG* protein and mRNA (Figure 7B & **Supplementary Figure S12C**). In strains with only a single CJnc190 promoter intact [C-190(P1) or C-190(P2)], target levels were intermediate between WT and Δ180/190, in line with the ∼2-fold difference in mature CJnc190 levels in these strains (**Supplementary Figure S12D**).

**Figure 7.**
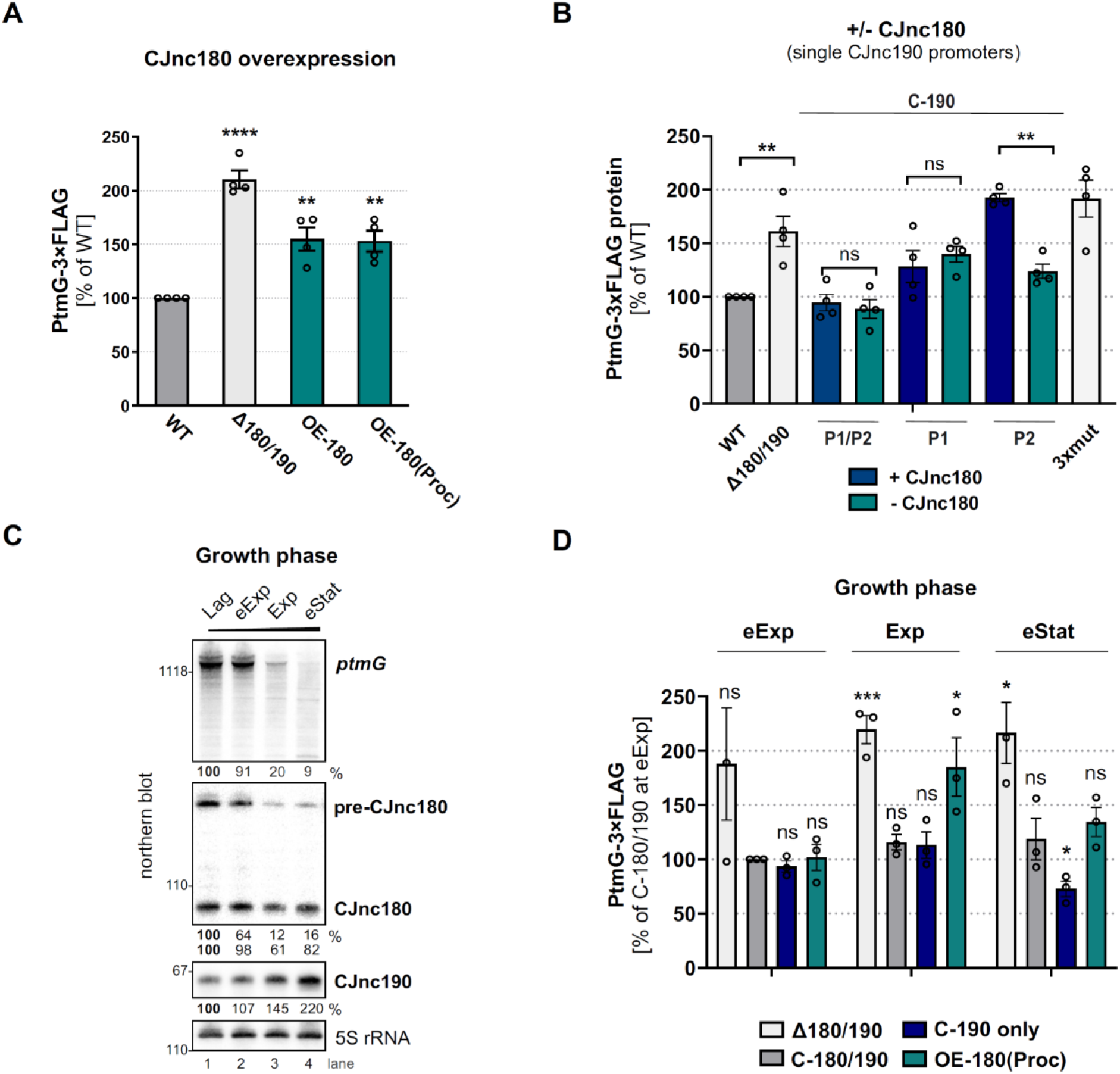
CJnc180 antagonizes CJnc190-mediated repression of *ptmG*. **(A)** The effect of CJnc180 overexpression on *ptmG*. OE-180: overexpression (second copy) of full-length CJnc180 from *rdxA*. OE-180(Proc): overexpression (second copy) of mature CJnc180 fused to the *E. coli rrnB* terminator (see Figure 1D). Levels of PtmG-3×FLAG protein were measured by western blot in the indicated strains in log phase. Error bars: SEM from four independent replicates. Student’s unpaired *t*-test vs. WT: **: p<0.01, ****: p<0.0001. See also **Supplementary Figure S12A. (B)** Absence of CJnc180 derepresses *ptmG* when CJnc190 is expressed from a single promoter. Levels of PtmG-3×FLAG protein were measured by western blot in the indicated strains in log phase. 3×mut: Δ180/190 complemented with CJnc180/190 carrying point mutations in all three validated promoters. Error bars: SEM from four independent replicates. Student’s unpaired *t*-test vs. WT: **: p<0.01, ns: not significant. See also **Supplementary Figure S12C & S12D. (C)** Northern blot analysis of precursor, mature sRNA, and *ptmG* target mRNA expression in WT at different growth phases in rich medium under microaerobic conditions. Lag: lag phase, eExp: early exponential, Exp: exponential, eStat: early stationary phase (OD_600_ 0.1, 0.25, 0.5, and 0.9, respectively). **(D)** The effect of the CJnc180 antagonist at different growth phases. Levels of PtmG-3×FLAG protein were measured in the indicated strains at three growth phases by western blot. Error bars: SEM from three independent replicates. Student’s unpaired *t*-test vs. WT: ***: p<0.001, *: p<0.05, ns: not significant vs. C-180/190 (dark grey bars) in eExp. See also **Supplementary Figure S13.** For all northern blots, probes for the mature sRNAs (CSO-0189 and CSO-0185 for CJnc180 and CJnc190, respectively) and the 5’ end of the *ptmG* ORF (CSO-1666) were used. As a loading control, 5S rRNA (CSO-0192) or RnpB (CSO-0497) was also probed.

We then wondered when CJnc180 antagonism might come into play. Specific conditions or factors regulating the sRNA pair are not yet known. However, examination of CJnc180 and CJnc190 levels in WT over growth showed that while mature CJnc190 levels increase, levels of CJnc180 fall or remain constant (precursor or mature, respectively) and *ptmG* mRNA levels decrease (Figure 7C). We determined the effect of CJnc180 absence/presence on PtmG-3×FLAG levels at different phases of growth (early exponential, mid exponential, and early stationary). We compared PtmG-3xFLAG protein and mRNA levels in Δ180/190, C-190-only, and OE-180(Proc) strains to those in C-180/190 as a control. PtmG-3×FLAG protein levels were relatively similar in the C-180/190 complemented strain at all culture densities, and in Δ180/190 showed sustained upregulation (Figure 7D). In contrast, CJnc180 absence or overexpression differentially affected PtmG::3×FLAG levels depending on growth phase. For C-190-only (CJnc180 absent, CJnc190 P1 and P2 active), PtmG-3×FLAG levels were mildly decreased compared to C-180/190 only in early stationary. In contrast, overexpression of CJnc180 de-repressed PtmG-3×FLAG levels in mid-log, but had no significant effect in stationary phase. Together, these experiments indicate that CJnc180 acts as a *cis*-acting sRNA antagonist of CJnc190, and that changing the ratios of the two sRNAs determines the outcome of *ptmG* regulation. Further analysis of CJnc190 expression from either promoter over growth did not reveal marked differences in expression (**Supplementary Figure S14A & S14B**). Moreover, we did not detect strong regulation of CJnc180 or CJnc190 promoter activity over growth (**Supplementary Figure S14C**). Therefore, the signals controlling the levels of the two sRNAs remain to be identified.

## DISCUSSION

Our study of the CJnc190/CJnc180 antisense sRNA pair provides insight into post-transcriptional regulation in the food-borne pathogen *C. jejuni*, as well as more generally into the complex cross-talk among RNA molecules and the role of RNase III in sRNA biogenesis. We have shown that the sRNAs from the virulence-associated CJnc180/190 locus of *C. jejuni* (Alzheimer et al., 2020) are processed using a complex biogenesis pathway involving RNase III (summarized in Figure 8). We demonstrated that one of these sRNAs, CJnc190, acts as a direct post-transcriptional repressor of the mRNA of the flagellin modification factor PtmG (Alzheimer et al., 2020; Howard et al., 2009), and that this regulation is antagonized by the *cis*-encoded CJnc180 sRNA. Although the sRNAs are expressed antisense to each other and both are processed by RNase III, only processing of CJnc180 requires its antisense partner, and CJnc190 is processed independently of CJnc180.

**Figure 8.**
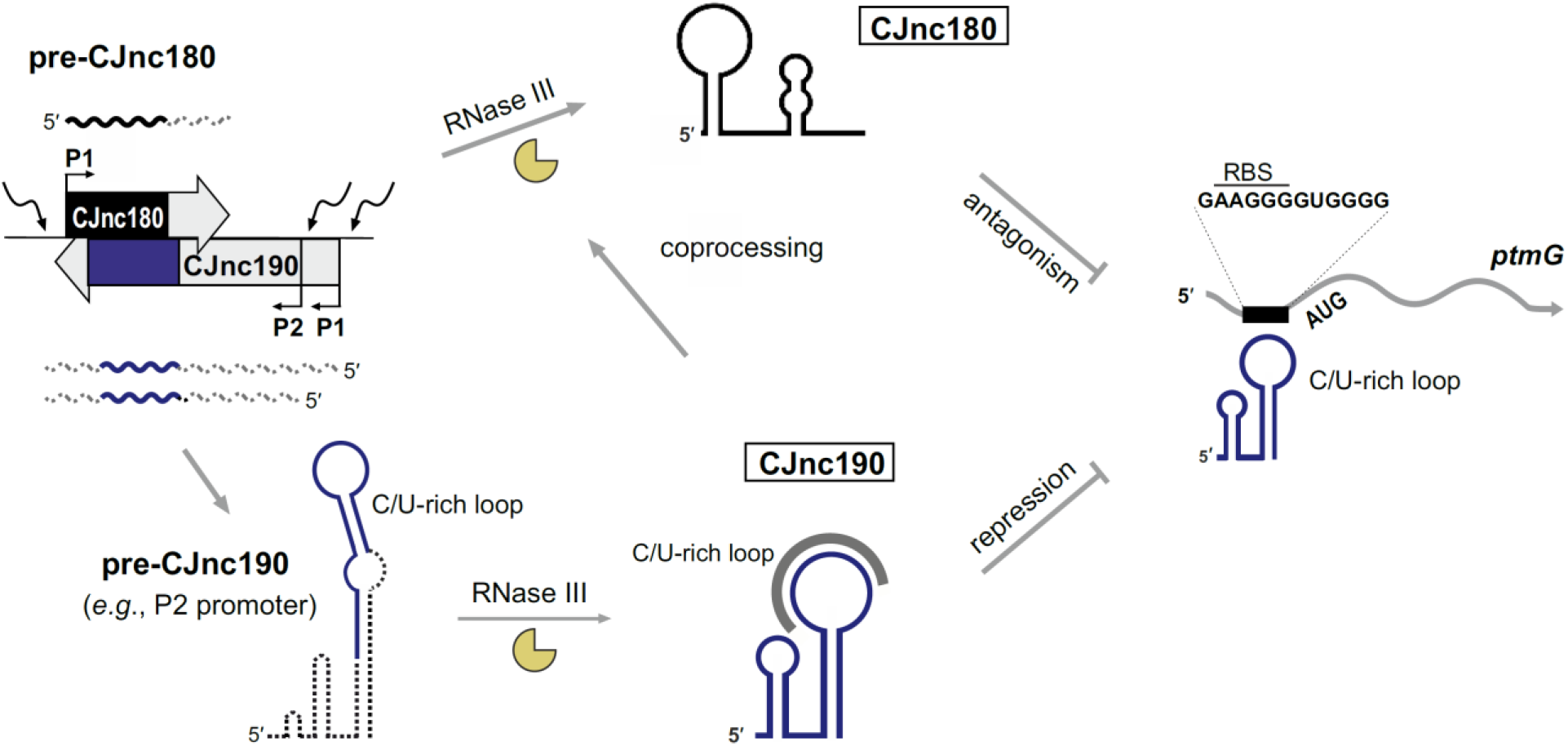
CJnc180/190 biogenesis, interplay, and regulation of *ptmG*. (Left) CJnc180/CJnc190 precursors are transcribed from one/two promoters, respectively, in response to so far still unknown signals/regulators. CJnc190 precursors harbour a long duplex structure involving regions flanking the mature Cjnc190 sRNA (blue). Mature CJnc190 is processed from the extended duplex structure in pre-CJnc190 by RNase III in the absence of CJnc180. In contrast, processing of pre-CJnc180 requires both RNase III and duplex formation with CJnc190. Mature CJnc190 represses translation of *ptmG* mRNA, encoding a colonization factor, by base-pairing between its C/U-rich loop and the G-rich *ptmG* RBS. Antisense CJnc180 antagonizes CJnc190 levels/activity by sequestration, decay, and/or transcriptional interference.

### Role of RNase III in sRNA biogenesis

While many enterobacterial sRNAs are generated or activated via processing by RNase E (Chao et al., 2017), our study revealed RNase III as a crucial factor for CJnc190 sRNA biogenesis in *C. jejuni*. Only a handful of RNase III-processed bacterial sRNAs have been described so far (Faubladier et al., 1990; Lalaouna et al., 2019). Our data establishes a role for RNase III in CJnc190 maturation, which in turn is required for *ptmG* regulation. However, it remains unclear why CJnc190 has this complex biogenesis pathway. Processing seems to generate a more stable form of CJnc190, but might also affect its activity. Surprisingly, CJnc190 processing was independent of its antisense RNA CJnc180. Instead, CJnc190 maturation involves co-processing on both sides of the mature sRNA by RNase III via a predicted long duplex region involving both ends of the precursor.

As it remains unclear how cleavage of the long CJnc190 5′/3′ end duplex might give rise to the final mature sRNA, additional RNases such as RNase J, Y, R or PNPase might be involved in further processing or trimming to the mature 5’/3’ ends. For example, in *E. coli*, processing of ribosomal RNAs or prophage-encoded DicF is initiated by RNase III cleavage of a stem-loop, followed by additional cleavages by RNases such as RNase E (Faubladier et al., 1990), and RNase III cleavage is followed by RNase J1 trimming during processing of the SRP RNA component scRNA (small cytoplasmic RNA) in *B. subtilis* (Yao et al., 2007).

In contrast to CJnc190, CJnc180 processing by RNase III required its *cis*-encoded partner to create a double-stranded substrate. The toxin mRNAs of several type I toxin-antitoxin (TA) systems have long been known to be processed by RNase III upon interaction with their antitoxin RNAs (Gerdes et al., 1992; Vogel et al., 2004), and in some bacteria such as *Bacillus* this even underlies RNase III essentiality (Durand et al., 2012). Besides processing of type I TA loci, RNase III has been implicated in degradation and processing of diverse sense/antisense RNA pairs, including those encoded on plasmids (Blomberg et al., 1990), such as antisense RNA mediated mRNA processing (Opdyke et al., 2011), as well as ribosomal RNA maturation (Iost et al., 2019), sRNA-mRNA target pair co-degradation (Afonyushkin et al., 2005; Romilly et al., 2012; Viegas et al., 2011), and CRISPR/tracrRNA co-processing in Cas9-based CRISPR/Cas systems (Deltcheva et al., 2011; Dugar et al., 2013).

### CJnc190 is antagonized by the *cis*-encoded CJnc180 sRNA

While CJnc180 is dispensable for CJnc190 processing and *ptmG* repression, we have shown *in vivo* that CJnc180 is a *cis*-acting antagonist of CJnc190, which can affect *ptmG* regulation. CJnc180 therefore appears to be a new *cis*-acting representative of sRNAs that target other sRNAs (reviewed in (Denham, 2020; Figueroa-Bossi and Bossi, 2018; Grüll and Massé, 2019)). Such antagonists are mainly *trans*-encoded and can be derived from mRNAs, tRNA 3’ external transcribed spacers or other sRNAs. In contrast to reported examples, CJnc180 antagonizes a *cis*-encoded sRNA. Additional candidate antagonizing RNAs are *cis*-encoded pairs that are differentially expressed under specific conditions (Denham, 2020). For example, antisense SraC/SdsR (RyeA/RyeB), widely conserved in *E. coli* and *Salmonella* (Fröhlich et al., 2016), show reciprocal expression and are also processed by RNase III (Vogel et al., 2003). In *E. coli*, SdsR overexpression leads to cell death via repression of *yhcB*, which is rescued by SraC overexpression, and the pair was also proposed as a novel toxin-antitoxin system where both components are sRNAs (Choi et al., 2019, 2018; Gupta et al., 2019). Whether the *cis*-acting antagonist CJnc180 from the CJnc190/180 locus also has an additional function and can act on other RNAs in *trans* remains to be seen.

While the exact mechanism of antagonism is unclear, hypotheses can be made based on other *cis*-acting sRNAs (Brantl, 2007). The two sRNAs could sequester each other, or play a role in each other’s turnover. CJnc180:CJnc190 co-processing would disrupt SL1 of CJnc190, which potentially protects the sRNA from RNase-mediated degradation. Antagonism could thus occur via cleavage and decay of CJnc190. Inspection of CJnc190 primer extension analysis for evidence of co-processing of the two sRNAs *in vivo* (**Supplementary Figure S8B**) did not reveal an RNase III- and CJnc180-dependent CJnc190 5′ end that would be consistent with the 2 nt 3′-overhangs generated by RNase III (Figure 6A, bottom), although reduced stability might preclude its detection. Alternatively, only the CJnc180 strand of the duplex might be cleaved, as has been reported for some RNase III substrates (Altuvia et al., 2018; Court et al., 2013; Dunn, 1976; Le Rhun et al., 2017).

Since overexpression of either “pre-processed” or full-length CJnc180 affected *ptmG*, co-processing might not be the only mechanism by which CJnc180 influences CJnc190. The extensive complementarity remaining for the processed sRNAs means that even mature CJnc180 could sequester CJnc190 - and possibly also promote degradation by RNase III. Finally, transcriptional interference might be involved, in line with our observation that abolishing CJnc180 expression reduced levels of some CJnc190 precursors with shorter 3′ ends (Figure 3D). The CJnc190 3′ end position is immediately upstream of the CJnc180 promoter on the opposite strand (**Supplementary Figure 3**). RNAs (including sRNAs, mRNAs, or derivatives of other cellular RNAs) that target sRNAs have commonly been termed “sponge” RNAs or competing endogenous RNAs (ceRNAs) (Denham, 2020; Grüll and Massé, 2019). Ultimately, the impact of CJnc180 on CJnc190 (and vice versa) might be multifactorial and include sequestration from targets, transcriptional interference, or promotion of decay. This might depend on their relative levels and/or on levels of other cellular molecules such as mRNA targets and RNases, as post-transcripional regulation is inherently dependent on the overall cellular state (Gottesman, 2004). We therefore propose terming CJnc180 (and also CJnc190) as RNA “antagonists”, and suggest reserving the term “sponge” for those that purely sequester or compete with targets without promoting decay.

The potential for independent regulation of CJnc180 and CJnc190 makes an antagonistic relationship attractive. The mechanism by which CJnc180 affects CJnc190 appears to be dependent on the stoichiometry of the two sRNAs, as has been previously proposed (Denham, 2020) and recently demonstrated for the RNAIII antagonist SprY (Le Huyen et al., 2021). A similar variation in sponge and sRNA stoichiometry has been reported for the ChiX sRNA and its decoy, the *chbBC* intergenic region (Figueroa-Bossi et al., 2009; Plumbridge et al., 2014). Presumably, CJnc180 could serve to buffer and/or set a threshold for CJnc190 levels to regulate targets, as for tRNA-derived sponge RNAs in *E. coli* (Lalaouna et al., 2015). While we observed that the mature sRNAs show inverse expression levels during growth, it remains unknown how they are transcriptionally controlled and in response to which signals their levels change. While CJnc180 is transcribed from one promoter (P1), CJnc190 is transcribed from at least two promoters (P1&P2). Precursors from both promoters are expressed during routine growth and give rise to the mature sRNA. The two promoters presumably increase the potential for environmental inputs into CJnc180/190, thereby increasing the complexity of the locus even further. Future work will reveal transcriptional regulators and conditions that regulate the CJnc180/190 promoters, and how transcriptional control intersects with RNase III processing.

### CJnc190 directly represses *ptmG* by targeting a G-rich sequence and impacts virulence

Based on *in vitro* and *in vivo* analyses, we demonstrated that CJnc190 directly represses *ptmG* translation by base-pairing with the G-rich RBS of its mRNA using a C/U-rich loop. Thus, CJnc190 binding likely interferes with translation initiation, resembling the canonical mode of sRNA-mediated target repression (Storz et al., 2011). The structure of mature CJnc190 is reminiscent of RepG sRNA from *H. pylori*, which uses a C/U-rich loop to target a phase-variable G-repeat in the 5’UTR of a chemotaxis receptor mRNA (Pernitzsch et al., 2014). Although CJnc190 and RepG share C/U-rich loop regions, their biogenesis is strikingly different: while RepG is transcribed as a separate standing gene corresponding to the mature sRNA (Pernitzsch et al., 2014), CJnc190 is transcribed opposite to another sRNA (CJnc180). Also in other species, several sRNAs with C/U-rich target interaction sites have been reported (Boisset et al., 2007; Bronesky et al., 2016; Heidrich et al., 2017; Schmidtke et al., 2013), suggesting that targeting of G-rich sequences might be a more widespread phenomenon.

A previous study of CJnc180/190 in our 3D model of the human intestine (Alzheimer et al., 2020), showed that CJnc180/190 as well as *ptmG,* the target of CJnc190, are involved in infection. PtmG is part of a six-gene cluster in the flagellin glycosylation island of *C. jejuni* NCTC11168 that has been associated with livestock strains (Champion et al., 2005; Mourkas et al., 2020) and is in the pathway generating legionaminic acid sugar precursors that decorate its flagellins (Howard et al., 2009; Zebian et al., 2016). As PtmG is required for chicken colonization (Howard et al., 2009), CJnc190 might impact virulence phenotypes via *ptmG* regulation. However, as CJnc190 is also found in strains that lack PtmG such as strain 81-176 (Dugar et al., 2013), it likely has additional targets that remain to be identified that could account for its infection phenotype (Alzheimer et al., 2020). In addition, antagonist CJnc180, which indirectly affects *ptmG* levels, might have an additional function as a *trans*-acting sRNA that might target other mRNAs encoding factors affecting virulence. Future studies will reveal the complete regulon of each sRNA and the contribution of their direct targets to virulence phenotypes.

## MATERIALS AND METHODS

### Bacterial strains and culture conditions

*C. jejuni* strains (**Supplementary File 1 - Table S2**) were routinely grown either on Müller-Hinton (MH) agar plates or with shaking at 140 rpm in *Brucella* broth (BB) at 37°C in a microaerobic atmosphere (10% CO_2_, 5% O_2_). All *C. jejuni* media was supplemented with 10 μg/ml vancomycin. Agar was also supplemented with marker-selective antibiotics [20 μg/ml chloramphenicol, 50 μg/ml kanamycin (Kan), 20 μg/ml gentamicin (Gm), or 250 μg/ml hygromycin B (Hyg)] where appropriate. *E. coli* strains were grown aerobically at 37°C in Luria-Bertani (LB) broth or on LB agar supplemented with the appropriate antibiotics for marker selection.

### General recombinant DNA techniques

Oligonucleotide primers for PCR, site-directed mutagenesis, Sanger sequencing, and northern blot probing are listed in (**Supplementary File 1- Table S3**) and were purchased from Sigma. Plasmids generated and/or used in this study are listed in (**Supplementary File 1 - Table S4**). Site-directed mutagenesis was performed on plasmids by inverse PCR with mutagenic primers as listed in (**Supplementary File 1 - Table S5**), according to standard protocols. DNA constructs and mutations were confirmed by Sanger sequencing (Macrogen). Restriction enzymes, *Taq* polymerase for validation PCR, and T4 DNA ligase were purchased from NEB. For cloning purposes, Phusion high-fidelity DNA polymerase (Thermo Fisher Scientific) was used. For PCR amplification of constructs containing the Hyg^R^ cassette, 3% DMSO was included in reactions.

### Transformation of *C. jejuni* for mutant construction

All *C. jejuni* mutant strains (deletion, chromosomal 3×FLAG-tagging, chromosomal point mutations, listed in (**Supplementary File 1 - Table S2**)) were constructed by double-crossover homologous recombination with DNA fragments introduced by electroporation or natural transformation. For electroporation, strains grown from frozen stocks until passage one or two on MH agar were harvested into cold electroporation solution (272 mM sucrose, 15% (v/v) glycerol) and washed twice with the same buffer. Cells (50 μl) were mixed with 200-400 ng PCR product on ice and electroporated (Bio-rad MicroPulser) in a 1 mm gap cuvette at 2.5 kV. Cells were then transferred with *Brucella* broth to a non-selective MH plate and recovered overnight at 37°C microaerobically before plating on the appropriate selective medium and incubating until colonies were visible (2-4 days). For natural transformation, approximately 100-1000 ng of genomic DNA isolated from the desired donor strain was mixed with the acceptor strain on non-selective MH plates and incubated for 4-5 h microaerobically at 37°C. The transformation mixture was then transformed to the appropriate selective medium and incubated until colonies were visible.

### *C. jejuni* deletion mutant construction by recombination with overlap PCR products

Deletion of the CJnc180/190 locus with a polar deletion cassette in *C. jejuni* NCTC11168 has been previously described (Alzheimer et al., 2020). Non-polar deletion mutants of protein-coding genes were constructed by homologous recombination with overlap PCR products consisting of a resistance cassette in between approximately 500 bp of sequence upstream and downstream of the target gene using primer pairs listed in (**Supplementary File 1- Table S6**). As an example, deletion of *rnc* in *C. jejuni* NCTC11168 with a non-polar Hyg^R^ cassette is described. The approximately 500 bp region upstream of *rnc* (Cj1635c) was amplified using CSO-0240/0241, while the downstream region was amplified using CSO-0242/0243. The 5’-ends of the antisense primer for the upstream region and the sense primer for the downstream region included regions overlapping the resistance cassette 5’- and 3’-end, respectively. The Hyg^R^ cassette was amplified using primers CSO-1678/1679 from pACH1 (Cameron and Gaynor, 2014). Next, a three-fragment overlap PCR was performed using the *rnc* upstream, *rnc* downstream, and Hyg^R^ cassette fragments in an equimolar ratio and primers CSO-0240/0243. Following confirmation of the correct size by gel electrophoresis, the resulting overlap PCR product was electroporated as described above into WT and deletion mutants were selected on plates containing hygromycin. The deletion strain (Δ*rnc*) was confirmed using a primer binding upstream of the *rnc* upstream fragment (CSO-0239) and an antisense primer binding the Hyg^R^ cassette (CSO-2857). For non-polar Kan^R^ and Gm^R^ deletions, cassettes were amplified using HPK1/HPK2 (Pernitzsch et al., 2014; Sharma et al., 2010) and pGG1 (Dugar et al., 2016) or pUC1813-apra (Bury-Moné et al., 2003) as a template, respectively.

### C-terminal 3×FLAG-tagging of *ptmG*

A 3×FLAG epitope was fused to the penultimate codon of *ptmG* at its native locus by homologous recombination with an overlap PCR product, which contained the upstream region of *ptmG* and its CDS to the penultimate codon fused to the 3×FLAG sequence, a Gm^R^ resistance cassette, and the *ptmG* downstream region. The *ptmG* upstream and coding region was amplified using CSO-1538/1539, and 500 bp of the downstream was amplified with CSO-1540/1541. A fusion of 3×FLAG to the Gm^R^ cassette was amplified using JVO-5142/HPK2 on the previously published 3×FLAG strain (*fliW*::3×FLAG)(Dugar et al., 2016).

### Heterologous expression from *rdxA*

The *rdxA* locus (Cj1066) can be used for heterologous gene expression in *C. jejuni* (Ribardo et al., 2010). Constructs for complementation in *rdxA* were made in plasmids containing approximately 500 bp of upstream and downstream sequence from *rdxA* flanking a Cm^R^ or Kan^R^ cassette (with promoter and terminator) by subcloning the *C. jejuni* sequence into previously-constructed plasmid vectors based on pST1.1 (Dugar et al., 2018) or pGD34.7 (Alzheimer et al., 2020). The CJnc180/190 and *ptmG* complementation plasmids (pGD34.7 and pSSv63.1) have been described previously (Alzheimer et al., 2020). Primers CSO-2276/2277 were then used to amplify all constructs from *rdxA*-based plasmids for electroporation into *C. jejuni*. Clones with intended insertions were validated by colony PCR using CSO-0643/0349 (*rdxA*-Cm^R^ constructs) or CSO-0023/0349 (*rdxA*-Kan^R^ constructs). Insertions were sequenced with CSO-3270, CSO-0643, or CSO-0023.

### Generation of plasmids for expression of processed/truncated (5′ or 3′) CJnc190 or CJnc180 (3′) at *rdxA*

To express 5′ end, pre-processed CJnc190 (CJnc190) from its native promoter P1, the 5’ end of CJnc190 precursors, up to the 5’ end of CJnc190 detected by primer extension, was removed from pGD34.7 by inverse PCR with primers CSO-2109/1545. After *Dpn*I digestion, ligation, and transformation into *E. coli*, the correct plasmid (pSSv20.1) was verified by colony PCR with pZE-A/pZE-XbaI and sequencing using CSO-0354. Truncations in the 5’ hairpin (sites A and A’) were generated in a similar fashion, using inverse PCR on pSSv80.1 with primers CSO-5235/1545 (A, pSSv148.1) or CSO-5234/1545 (A’, pSSv147.1). Following *Dpn*I digestion, ligation, and transformation into *E. coli* TOP10, positive clones were identified by colony PCR with CSO-0643/3270 and sequenced with CSO-0643. All CJnc190 3’ end truncations were generated by inverse PCR with CSO-0347/5391 on plasmids pSSv20.1, pSSv147.1, pSSv148.1, and pSSv80.1 to generate plasmids pSSv160.2, pSSv162.1, pSSv163.1, and pSSv161.1, respectively. Clones were also identified by colony PCR with CSO-0643/3270 and sequenced with CSO-0643. For the expression of processed CJnc180 with an *E. coli rrnB* terminator, plasmid pGD34.7 was used as template for inverse PCR using the primers CSO-3831/3832. After *Dpn*I digestion, ligation, and transformation into *E. coli*, the correct plasmid (pSSv116.1) was verified by colony PCR with CSO-0643/3270 and sequencing using CSO-3270.

### Generation of a plasmid for expression from the Cj0046 pseudogene locus

The Cj0046 pseudogene locus has been used previously as a *C. jejuni* heterologous expression/complementation locus (Kim et al., 2008). To generate a plasmid allowing insertion of genes with a resistance cassette within Cj0046, ∼1000 bp of the Cj0046 locus was first amplified using primers CSO-1402/1405. Recipient plasmid pSSv1.2 (CSS-1125 (Dugar et al., 2016)) was amplified by inverse PCR with CSO-0073/0075. The vector and insert were digested with *Xba*I and *Xho*I and ligated together. A positive plasmid clone (pSSv53.1, CSS-2861) was validated by colony PCR with CSO-1402/1405 and sequencing with CSO-1402. Next, pSSv53.1 was amplified by inverse PCR with CSO-2748/2751 and digested with *Eco*RI. A polar (promoter and terminator) Gm^R^ cassette was amplified with CSO-2749/2750 on pGD46.1 (CSS-0858) and similarly digested. The pSSv53.1 backbone and Gm^R^ cassette were ligated together to generate pSSv54.3, which was validated by colony PCR with CSO-1402/1405.

### Generation of a plasmid for insertion of CJnc190 at Cj0046

To generate plasmids containing CJnc180/190 alleles with a Gm^R^ cassette flanked by 500 bp of Cj0046, the WT sRNA region was first amplified from *C. jejuni* NCTC11168 WT (CSS-0032) gDNA with CSO-0354/0355. The pSSv54.3 plasmid (CSS-2872) was then amplified by inverse PCR with CSO-2750/2751. Both insert and vector were digested with *Nde*I and *Cla*I and ligated. A positive clone (pSSv54.3, CSS-2872) was identified by colony PCR with CSO-0833/0354 to generate plasmid pSSv55.4. Point mutant alleles were generated by site-directed mutagenesis with primer pairs listed in (**Supplementary File 1- Table S5**). All mutations were validated by sequencing with CSO-0833. For insertion of processed CJnc190 into the Cj0046 plasmid (pSSv56.6), the sRNA region from pSSv20.1 (CSS-1610) was amplified by PCR with CSO-2839/0354 and the pSSv54.3 (CSS-2872) vector was amplified with CSO-2750/2751. Both vector and insert were digested with *Nde*I and *Cla*I and ligated. A positive clone was identified by colony PCR with CSO-2839/1405 and sequencing with CSO-2839.

### Construction of *ptmG(10th)*-GFP translational fusions at Cj0046 for point mutant analysis

A *gfpmut3* translational fusion reporter for *ptmG* was constructed as follows, starting with plasmid pST1.1(Dugar et al., 2018). First, the *metK* promoter of pST1.1 was replaced with the promoter, 5’ UTR, and the first ten codons of *ptmG*. The *ptmG* 5’ UTR and first ten codons were amplified by PCR using CSO-1670/1671 and digested with *Nde*I. Plasmid pST1.1 was then amplified by PCR with primers designed to remove the *metK* promoter (CSO-1669/0762) and similarly digested. Ligation of the pST1.1 backbone and *ptmG* region resulted in plasmid pSSv15.1, which was confirmed by colony PCR with CSO-1670/0348 and sequencing with CSO-0023. Next, the *ptmG(10th)*-GFP reporter fusion was moved into the Cj0046 pseudogene insertion plasmid pSSv54.3 as follows. The region of pSSv15.1 harbouring the reporter and Kan^R^ cassette was amplified using CSO-0159/2877, and pSSv54.3 was amplified using CSO-2818/2870, which removes the Gm^R^ cassette. Both insert and vector were digested with *Pst*I and *Not*I and then ligated to make pSSv61.1, which was validated by colony PCR with CSO-0023/0789. A single G-C point mutation (M1) was introduced into the 5’ UTR of *ptmG* in pSSv61.1 by inverse PCR with the mutagenic primer pair CSO-2875/2876 to create pSSv62.1. The mutation was confirmed by sequencing with CSO-0789. Next, the *rdxA*::Kan^R^-*ptmG*UTR-GFP construct from the WT and M1 plasmids was amplified using CSO-1402/1405 and transformed into *C. jejuni*. Insertions were confirmed by colony PCR with CSO-0023/3217 and sequencing with CSO-0023. Strains in these backgrounds deleted for the sRNAs and/or complemented at *rdxA* were then constructed as for the WT background, described above, except for the addition of CJnc190 M1 point mutant, which was made by site-directed mutagenesis of pSSv20.1 with CSO-2871/2872 and confirmed by sequencing with CSO-0643.

### Generation of promoter exchange experiment strains

The *ptmG* promoter (P*_ptmG_*) was exchanged for the *flaA* promoter (P*_flaA_*) in the above *ptmG(10th)*-GFP translational fusion as follows. Plasmid pSSv61.1 (CSS-2921) was amplified by inverse PCR with CSO-0762/1955 and the *flaA* promoter was amplified from WT gDNA (CSS-0032) with CSO-1732/1733. Both amplicons were digested with *Xma*I and ligated. A positive clone (pSSv83.1, CSS-3282) was validated by colony PCR with CSO-0023/1933 and sequencing with CSO-0023. The region of interest was amplified from pSSv83.1 with primers CSO-1402/1405 and transformed into *C. jejuni* Δ*ptmG*_UTR (CSS-3234). Positive clones were checked by colony PCR with CSO-0023/0789 to validate strain CSS-3302.

A control fusion of the *flaA* promoter was constructed as follows. First, the intermediate plasmid pGD7.1 (Alzheimer et al., 2020) was amplified by inverse PCR with CSO-0482/0493, digested with *Not*I/*Pst*I, and ligated with a similarly digested polar Kan^R^ cassette-P*_ureA_*-*gfpmut3* PCR product amplified with CSO-0513/0414 from p463 (CSS-0079, (Pernitzsch et al., 2014) to generate plasmid pMA4.5 (CSS-2389), which was validated by colony PCR with CSO-0348/0513 and sequencing with CSO-0345/0348. The backbone of pMA4.5 was then amplified by inverse PCR with CSO-1734/0762, and the *flaA* (Cj1339c) promoter was amplified from *C. jejuni* NCTC11168 WT (CSS-0032) gDNA with CSO-1732/1733. Both vector and insert were digested with *Xma*I and *Eco*RI and ligated to create pST10.1 (CSS-2379), which was validated by colony PCR with CSO-0023/0348 and sequencing with CSO-0348. The reporter regions were amplified with CSO-2276/2277 for electroporation into *C. jejuni*.

The *fliA* gene (Cj0061) was disrupted in the pSSv61.1 (P*_ptmG_*-*ptmG(10th)*-GFP), pSSv83.1 (P*_flaA_*-*ptmG(10th)*-GFP), and pST10.1 (P*_flaA_*-*flaA*-GFP) promoter fusion strains (CSS-2945, CSS-3301, and CSS-3388, respectively) by natural transformation of genomic DNA from a strain carrying the *fliA* region with a non-polar gentamicin resistance cassette (CSS-1133, (Dugar et al., 2016)) to generate CSS-3983, CSS-3303, and CSS-4025. Deletion of *fliA* was confirmed by colony PCR with CSO-1153/HPK2. The CJnc180/190 region was disrupted in CSS-3301 and CSS-3388 with a polar gentamicin resistance cassette by transformation with an overlap PCR product generated as described above and validation by colony PCR with CSO-0246/HPK1 to generate strains CSS-3305 and CSS-3390. The P*_flaA_*-*ptmG(10th)*-GFP Δ180/190 strain (CSS-3305) was then complemented at *rdxA* with the mature CJnc190 region from pSSv20.1 to generate CSS-2957. For this, a PCR product amplified with CSO-2276/2277 on pSSv20.1 (CSS-1610) was electroporated into CSS-3305 and colonies were checked by colony PCR with CSO-0349/0643 and sequencing with CSO-0643.

### Generation of promoter point mutant alleles

For construction of different CJnc180/190 alleles, the original pGD34.7 or pSSv55.1 plasmids were subjected to site-directed mutagenesis by inverse PCR/*Dpn*I digestion using primers listed in (**Supplementary File 1 - Table S5**) to create otherwise isogenic plasmids listed in (**Supplementary File 1 - Table S4**). First, the CJnc180(P1) promoter was disrupted (see **Supplementary File 1 - Table S5** for mutations) to create strain C-190-only [CJnc190(P1) and CJnc190(P2) active] (plasmid pSSv38). To this allele, CJnc190(P1) or (P2) promoter inactivation mutations were added to create strains C-190(P2) only and C-190(P1) only (pSSv98 and pSSv97), respectively. Finally, a strain with all three putative promoters mutated (C-3×mut) was also constructed (pSSv79). The different *rdxA*::Cm^R^-CJnc180/190 alleles were amplified from the plasmids using CSO-2276/2277 and electroporated into the *C. jejuni* Δ180/190 strain (CSS-1157). The strains were validated for the correct insertion of each transformed allele by colony PCR using CSO-0643/0349 and sequencing with CSO-0643.

### Transcriptional fusions to superfolder GFP

The CJnc180 promoter from *C. jejuni* and CJnc180 upstream region from *C. coli* were fused to a promoterless superfolder GFP (sfGFP) cassette with the RBS from *hupB* (Cj0913c) by overlap PCR as follows. An *rdxA*UP(approx 500 bp)-Kan^R^-sfGFP cassette was first generated in a plasmid by amplification of sfGFP from pXG10 (Corcoran et al., 2012) with primers CSO-3279/3569, digestion with *Xma*I, and ligation to pST1 (Dugar et al., 2018). The pST1 backbone was amplified with CSO-0762/0347 and similarly digested with *Xma*I. This plasmid was validated by colony PCR with CSO-0023/3527 and sequencing with CSO-0023. The *rdxA*UP-Kan^R^-sfGFP cassette was then amplified with primers CSO-5590/2276, and the *rdxA*DN region (approx 500 bp) was amplified with CSO-0347/2277. CJnc180 promoter/upstream regions were amplified with CSO-5595/5593 and CSO-5597/5598 (*C. jejuni* and *C. coli*, respectively) from WT genomic DNA to introduce regions overlapping the *rdxA*DN fragment or *hupB* RBS. The three fragments (*rdxA*DN/promoter/sfGFP-Kan^R^-*rdxA*UP) were mixed, annealed, and amplified by overlap PCR with CSO-2276/2277. The resulting PCR product was electroporated into *C. jejuni* NCTC11168 WT. Kan^R^ colonies were validated for insertion of the transcriptional fusion at *rdxA* by colony PCR with CSO-0349/0789 and promoter regions were checked by sequencing with CSO-3270.

### Total RNA extraction and analysis by northern blotting

For analysis of total RNA, bacterial strains were grown to log phase in BB and approximately 4 OD_600_ were harvested and mixed with 0.2 volumes of stop-mix (95% ethanol and 5% phenol, v/v). Samples were immediately snap-frozen in liquid nitrogen and stored at −80°C until RNA extraction. Frozen samples were thawed on ice and centrifuged at 4°C to collect cell pellets (4,500 *g*, 20 min), which were then lysed by resuspension in 600 μl of a solution containing 0.5 mg/ml lysozyme and 1% SDS in Tris-EDTA buffer (pH 8.0) and incubation for 2 min at 64°C. Total RNA was extracted from the lysate using the hot-phenol method as described previously (Sharma et al., 2010). For northern blot analysis, 5 - 10 μg of total RNA in Gel Loading Buffer II (GLII, Ambion) was loaded per lane on 6% polyacrylamide (PAA) / 7 M urea denaturing gels in 1× TBE buffer. Following electrophoretic separation, RNA was transferred to Hybond-XL membranes (GE Healthcare) by electroblotting. Transferred RNA was then cross-linked to the membrane with ultraviolet light to the membrane and hybridized with γ^32^P-ATP end-labelled DNA oligonucleotides (**Supplementary File 1; Table S3**) in Roti Hybri-quick (Roth) at 42 °C overnight. Membranes were then washed 20 minutes each at 42° C in 5×, 1×, and 0.5× SSC (saline-sodium citrate) + 0.1% SDS, dried, and exposed to a phosphorimager screen. Screens were scanned using a FLA-3000 Series Phosphorimager (Fuji) and bands were quantified using AIDA software (Raytest, Germany).

### Total protein sample analysis by SDS-PAGE and western blotting

Analysis of protein expression in *C. jejuni* was performed by SDS-PAGE and western blotting. Bacterial cells were collected from cultures in mid-log phase (OD_600_ 0.4-0.5) by centrifugation at 11,000 *g* for 3 min. Cell pellets were resuspended in 100 μl of 1× protein loading buffer (62.5 mM Tris-HCl, pH 6.8, 100 mM DTT, 10% (v/v) glycerol, 2% (w/v) SDS, 0.01% (w/v) bromophenol blue) and boiled for 8 min. For analysis of total proteins, 0.05-0.1 OD_600_ of cells were loaded per lane on a 12% SDS-polyacrylamide gels. Gels were stained with PageBlue (Thermo Fisher Scientific, #24620). For western blot analysis, samples corresponding to an OD_600_ of 0.05-0.1 were separated on 12% SDS-PAA gels and transferred to a nitrocellulose membrane by semidry blotting. Membranes were blocked for 1 h with 10% (w/v) milk powder in TBS-T (Tris-buffered saline-Tween-20) and then incubated overnight with primary antibody (monoclonal anti-FLAG, 1:1,000; Sigma-Aldrich, #F1804-1MG; or anti-GFP, 1:1000, Roche #11814460001 in 3% BSA/TBS-T) at 4°C. Membranes were then washed with TBS-T, followed by 1 h incubation with secondary antibody (anti-mouse IgG, 1:10,000 in 3% BSA/TBS-T; GE-Healthcare, #RPN4201). All antibodies were dissolved in 3% BSA (bovine serum albumin)/TBS-T. After washing, the blot was developed using enhanced chemiluminescence reagent and imaged using an ImageQuant LAS-4000 imager (GE). Bands were quantified using AIDA software. As a loading control, a monoclonal antibody specific for GroEL (1:10,000; Sigma-Aldrich, #G6532-5ML) with an anti-rabbit IgG (1:10,000; GE Healthcare, #RPN4301) secondary antibody was used to probe membranes after FLAG/GFP.

### Rifampicin RNA stability assays

To determine the stability of CJnc180 and CJnc190 in *C. jejuni* NCTC11168 WT and Δ*rnc*, strains were grown to an OD_600_ of 0.5 (mid-log phase) and treated with rifampicin to a final concentration 500 μg/ml. Samples were harvested for RNA isolation as described above at indicated time points following rifampicin addition (2, 4, 8, 16, 32, 64 min). After RNA isolation, 10 μg of each RNA sample (DNase I-digested) was used for northern blot analysis as detailed above.

### Primer extension analysis of RNA 5′ ends

Total RNA was extracted from bacteria in log phase as described above. RNA was digested with DNase I (Thermo Fisher Scientific) to remove DNA, and then 5 - 10 μg of RNA was added to a total volume of 5.5 μl with H_2_O, denatured, and snap-cooled on ice. A 5’ end ^32^P labeled DNA oligonucleotide complementary to the RNA of interest was then added (**Supplementary File 1 - Table S3**) and annealed by heating to 80°C, followed by slow cooling (1°C per min) to 42°C. A master mix with reverse transcriptase (RT) buffer and 20 U Maxima RT (Thermo Fisher Scientific) was added and the reaction was allowed to proceed for 1 h at 50°C. Reactions were stopped with 12 μl GLII (Ambion, 95% (v/v) formamide, 18 mM EDTA, and 0.025% (w/v) SDS, xylene cyanol, and bromophenol blue). A sequencing ladder was also constructed using the DNA Cycle sequencing kit (Jena Bioscience) according to the manufacturer’s instructions with the CJnc180/190 region amplified with primers CSO-0354/0355 from genomic DNA (NCTC11168 wild-type) as template and the same radioactively-labeled primer was used for the reverse transcription reaction. Reactions were separated on 6 or 10% PAA-urea sequencing gels, which were then dried and exposed to a phosphorimager screen, and then scanned (FLA-3000 Series, Fuji). The following primers were used for primer extension: CJnc190 - CSO-0185; CJnc180 - CSO-0188.

### RACE analysis of RNA 3’-ends

Total RNA from *C. jejuni* WT grown to mid-log phase was used for RACE analysis of the 3’-ends of each sRNA. Briefly, 2 μl of 10× Antarctic Phosphatase buffer (NEB), 1 U Antarctic Phosphatase, and 10 U SUPERase•In RNase inhibitor was added to 7.5 μg of denatured/snap-cooled RNA in a 20 μl reaction and incubated for 1 h at 37°C. The reaction was then made up to 100 ul and extracted with an equal volume of phenol:chloroform:isoamyl alcohol (PCI) in a Phase-Lock gel tube (5PRIME). RNA was then precipitated with 7.5 μg GlycoBlue (Ambion) and 2.5 vol 30:1 Mix. The RNA was dissolved in water, 250 pmol RNA adaptor E1 (CSO-4916) was added, and the mixture was denatured and snap-cooled. To this was added 2 μl DMSO, 2 μl 10× T4 RNA ligase buffer (NEB), 20 U T4 RNA ligase enzyme, and 10 U SUPERase•In. Following ligation overnight at 16°C, the RNA was extracted with PCI and precipitated as described above. The RNA was then dissolved in water, denatured and snap-cooled, and subjected to reverse transcription with Maxima reverse transcriptase and DNA oligo E4 for 5 min at 50°C, 1 h at 55°C, and 15 min at 70°C. RNA was then removed by digestion with 5 U RNase H for 22 min at 37°C. Two microlitres of this reaction was then used as template for PCR using *Taq* DNA polymerase (NEB), DNA oligo E4 (CSO-4720), and either CSO-1380 (CJnc190) or CSO-1973 (CJnc180). Cycling conditions were as follows: 5 min at 95°C, 35 cycles of 95°C for 30 s - 57°C for 30 s - 72°C for 45 s, and 10 min at 72°C. Amplification was checked on a 2% agarose/1× TAE gel, reactions were cleaned up with the NucleoSpin Gel and PCR Clean-up Kit (Macherey-Nagel), and ligated to pGEM-T Easy (Promega) according to the manufacturer’s instructions. For CJnc180, the inserts of 9 white clones were sequenced, and for CJnc190, the inserts of ten white clones were sequenced with primer REV or UNI61.

### *In vitro* transcription and 5′ end-labeling of RNAs

PCR with Phusion DNA polymerase was used to generate DNA templates containing the T7 promoter sequence using oligonucleotides and DNA templates listed in **Supplementary File 1 - Table S7**. Transcription of RNAs *in vitro* by T7 RNA polymerase was then carried out using the MEGAscript T7 kit (Ambion) according to the manufacturer’s instructions. RNAs were then checked for quality by electrophoresis on a PAA-urea gel, dephosphorylated with Antarctic Phosphatase (NEB), 5′-end-labelled (γ^32^P) with polynucleotide kinase (ThermoFisher Scientific), and purified by gel extraction as previously described (Papenfort et al., 2006). Expected sequences of the resulting T7 transcripts are listed in **Supplementary File 1 - Table S7**.

### Electrophoretic mobility shift assays (EMSAs)

Gel shift assays were performed as described previously (Pernitzsch et al., 2014). Briefly, 5′ end radiolabeled RNA (0.04 pmol) was denatured (1 min, 95°C) and cooled for 5 min on ice. Yeast tRNA (1 μg, Ambion) and 1 μl of 10× RNA Structure Buffer (Final concentration 10 mM Tris, pH 7, 100 mM KCl, 10 mM MgCl_2_) was then mixed with the labeled RNA. Unlabeled RNA (2 μl diluted in 1× Structure Buffer) was added to the desired final concentrations (0 nM, 10 nM, 20 nM, 50 nM, 100 nM, 200 nM, 500 nM, or 1 μM). Binding reactions were incubated at 37°C for 15 min. Before loading on a pre-cooled native 6% PAA, 0.5× TBE gel, samples were mixed with 3 μl native loading buffer [50% (v/v) glycerol, 0.5× TBE, 0.2% (w/v) bromophenol blue]. Gels were run in 0.5× TBE buffer at 300 V and 4°C. Gels were dried, exposed to a phosphorimager screen, and then scanned (FLA-3000 Series, Fuji).

### Inline probing

Inline probing assays for RNA structure and binding interactions *in vitro* were performed essentially as described previously (Pernitzsch et al., 2014). End-labeled RNAs (0.2 pmol, see above) in 5 μl water were mixed with an equal volume of 2× Inline buffer (100 mM Tris-HCl, pH 8.3, 40 mM MgCl_2_, and 200 mM KCl and incubated for 40 h at room temperature to allow spontaneous cleavage. Reactions were stopped with an equal volume of 2× Colourless loading buffer (10 M urea and 1.5 mM EDTA, pH 8.0). Reactions were separated on 6 or 10% PAA-urea sequencing gels, which were dried and exposed to a PhosphorImager screen. RNA ladders were prepared using Alkaline hydrolysis buffer (OH ladder) or Sequencing buffer (T1 ladder) according to the manufacturer’s instructions (Ambion).

### RNase III cleavage assays

*In vitro*-transcribed pre-CJnc180 was 5′ end labeled as described for Inline probing and EMSA and subjected to RNase III cleavage assays as follows. Labeled pre-CJnc180 (0.2 pmol) was briefly denatured and snap-cooled on ice, followed by the addition of Structure buffer to a final concentration of 1× and yeast tRNA to a final concentration of 0.1 mg/ml. Where necessary, unlabeled mature CJnc190 (0.2 or 2 pmol) was denatured and snap-cooled separately and added to reactions. Reactions were pre-incubated at 37°C for 10 minutes, followed by the addition of RNase III (NEB; 1/625 U) and further incubation at 37°C for 5 minutes to allow limited cleavage. Reactions were stopped by the addition of an equal volume of GLII and separated on a 10% PAA-urea sequencing gel, which was then dried and exposed to a PhosphorImager screen. RNA ladders were prepared using Alkaline hydrolysis buffer (OH ladder) or Sequencing buffer (T1 ladder) according to the manufacturer’s instructions (Ambion).

### *In vitro* translation

*In vitro* translation of target mRNA reporter fusions in the presence and absence of sRNAs was carried out using the PURExpress system (NEB). An *in vitro*-transcribed RNA including the *ptmG* 5′ leader (including RBS, first ten codons, and CJnc190 binding site) and first 10 codons fused to *gfpmut3* (*ptmG(10th)-gfp*) was used as template for translation (**Supplementary File 1 - Table S7**). For each reaction, 4 pmol of denatured template RNA was incubated either alone or with equimolar (1×), 10×, or 50× ratios of sRNA species for 10 minutes at 37°C. *In vitro* translation components were then added, and reactions were incubated a further 2 hours at 37°C. Reactions were stopped with an equal volume of 2× protein loading buffer. One half of the reaction was analyzed by western blotting on 12% SDS-PAA gels with an antibody against GFP, and the second half was loaded on a second gel which was stained with PageBlue after electrophoresis, as a loading control.

### Differential RNA-seq data

Processed primary transcriptome data generated by dRNA-seq for *C. jejuni* NCTC11168 (Dugar et al., 2013) was retrieved from the NCBI Gene Expression Omnibus (GEO) using the accession GSE38883, was inspected using the Integrated Genome Browser (bioviz.org) (Freese et al., 2016).

### Mass spectrometry

Potential targets of CJnc180/190 were identified by mass spectrometry (MS/MS) analysis of gel-excised, trypsinized protein bands, performed by the Bioanalytical Mass Spectrometry lab at the Max-Planck-Institute for Biophysical Chemistry (https://www.mpibpc.mpg.de/urlaub) according to published standard protocols. Proteins were separated by SDS-PAGE (12% PAA) and stained with PageBlue protein staining solution (ThermoFisher Scientific) before analysis.

## ACKNOWLEDGEMENTS

This work was supported through a grant within the Bavarian Research Network bayresq.net (to C.M.S.). We are grateful to Uwe Plessmann and Prof. Dr. Henning Urlaub for acquiring and analyzing mass spectrometry data as well as Philipp Kible for technical assistance. We thank Isabelle Iost and Fabien Darfeuille for feedback on RNase III assays as well as Sharma lab members for critical comments on the manuscript.

## AUTHOR CONTRIBUTIONS

SLS & CMS designed the experiments; SLS performed the experiments; SLS & CMS analyzed data; SLS & CMS wrote the manuscript.

