## Supplemental Materials for "RNase III-mediated processing of a *trans*-acting bacterial sRNA and its *cis*-encoded antagonist"

**This file contains:**

**Supplementary Figures S1 – S14**

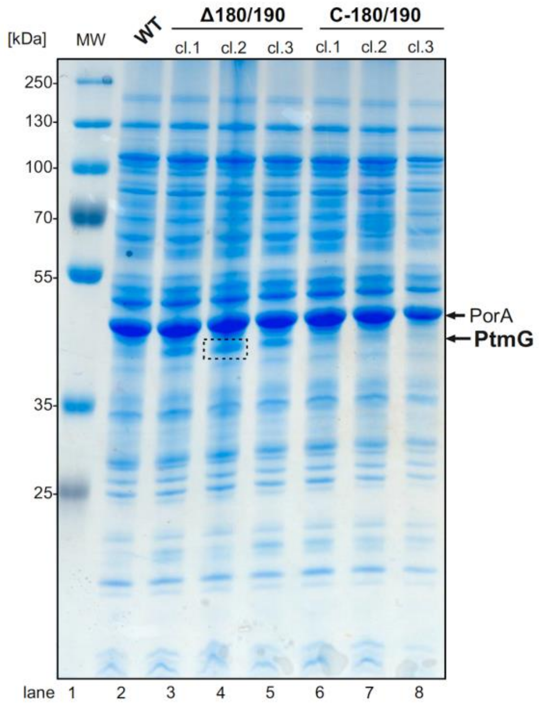


**Supplementary Figure S1. Identification of CJnc180/190 targets by SDS-PAGE and mass spectrometry analysis.** Samples were harvested for total protein analysis from the wild-type (WT) strain and three independent clones (cl.) of Δ180/190 and C-180/190 in mid-exponential phase ($OD_{600}$ approx 0.5). Proteins from 0.1 $OD_{600}$ of cells were separated per well on a 12% polyacrylamide SDS-PAGE gel and stained with PageBlue. The region surrounded by the dashed box was excised and subjected to trypsin digestion and mass spectrometry for identification of *C. jejuni* proteins (see **Supplementary File 1 - Table S1**). PorA served as a loading control. MW indicates a prestained protein molecular weight ladder.

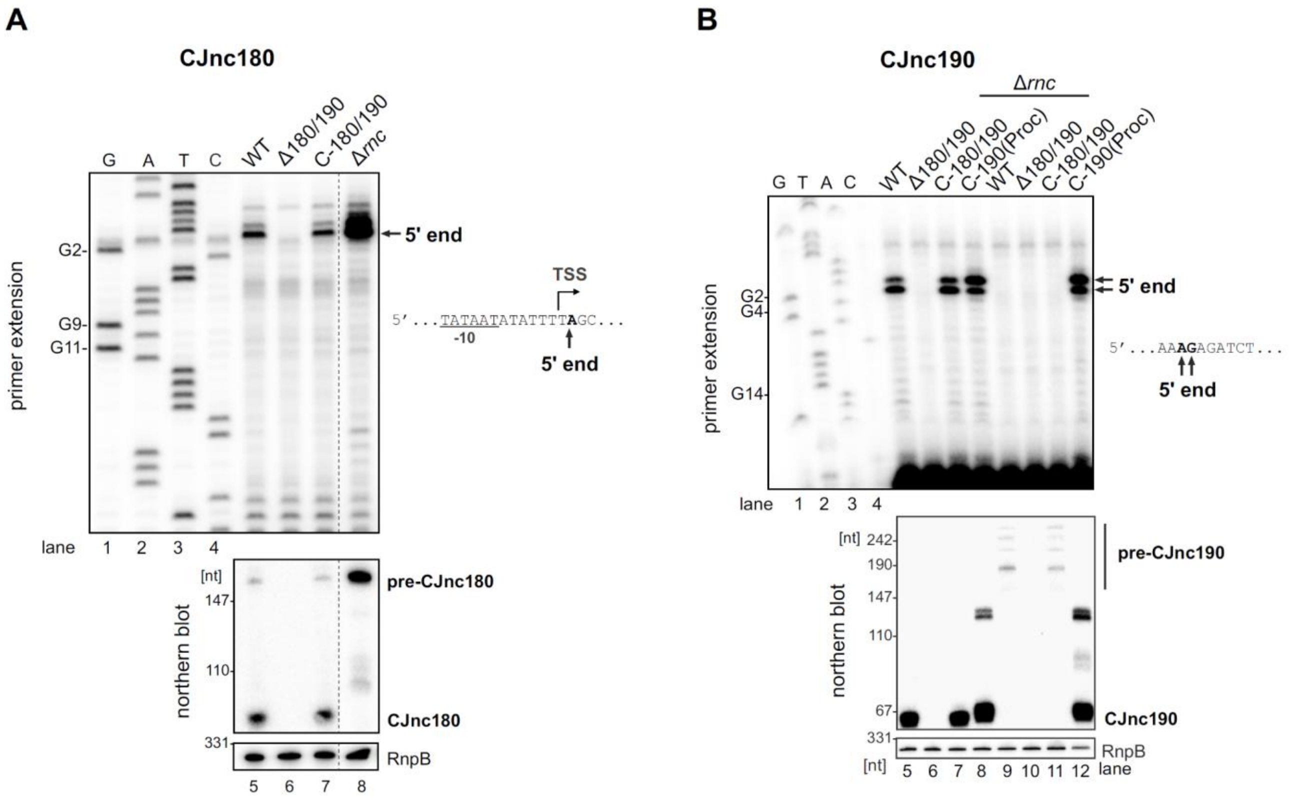


**Supplementary Figure S2. Mapping 5′ boundaries of mature CJnc180 and CJnc190 by primer extension. (A)** Total RNA from *C. jejuni* NCTC11168 WT, Δ180/190, Δ180/190 complemented with CJnc180/190, or Δ*rnc* was subjected to primer extension analysis with a probe annealing to mature CJnc180 (CSO-0189). **(B)** Total RNA from WT, Δ180/190, and Δ180/190 complemented with CJnc180/190 or CJnc190(Proc) (see main **Figure 1D**) was subjected to primer extension analysis with a probe annealing to mature CJnc190 (CSO-0185). The arrows mark detected 5' ends, which are also displayed in relation to the annotated TSS (Dugar et al., 2013) and promoter -10 box to the right. Lanes 1-4: CJnc180/190 sequencing ladders generated with the same probe. Northern blots of the same RNA were probed with the same oligonucleotides that are complementary to the mature sRNAs (CSO-0189/CSO-0185 for CJnc180/190) and RnpB as a loading control (CSO-0497). For panel A, intervening lanes were cropped from the blot/gel pictures (vertical dashed line).

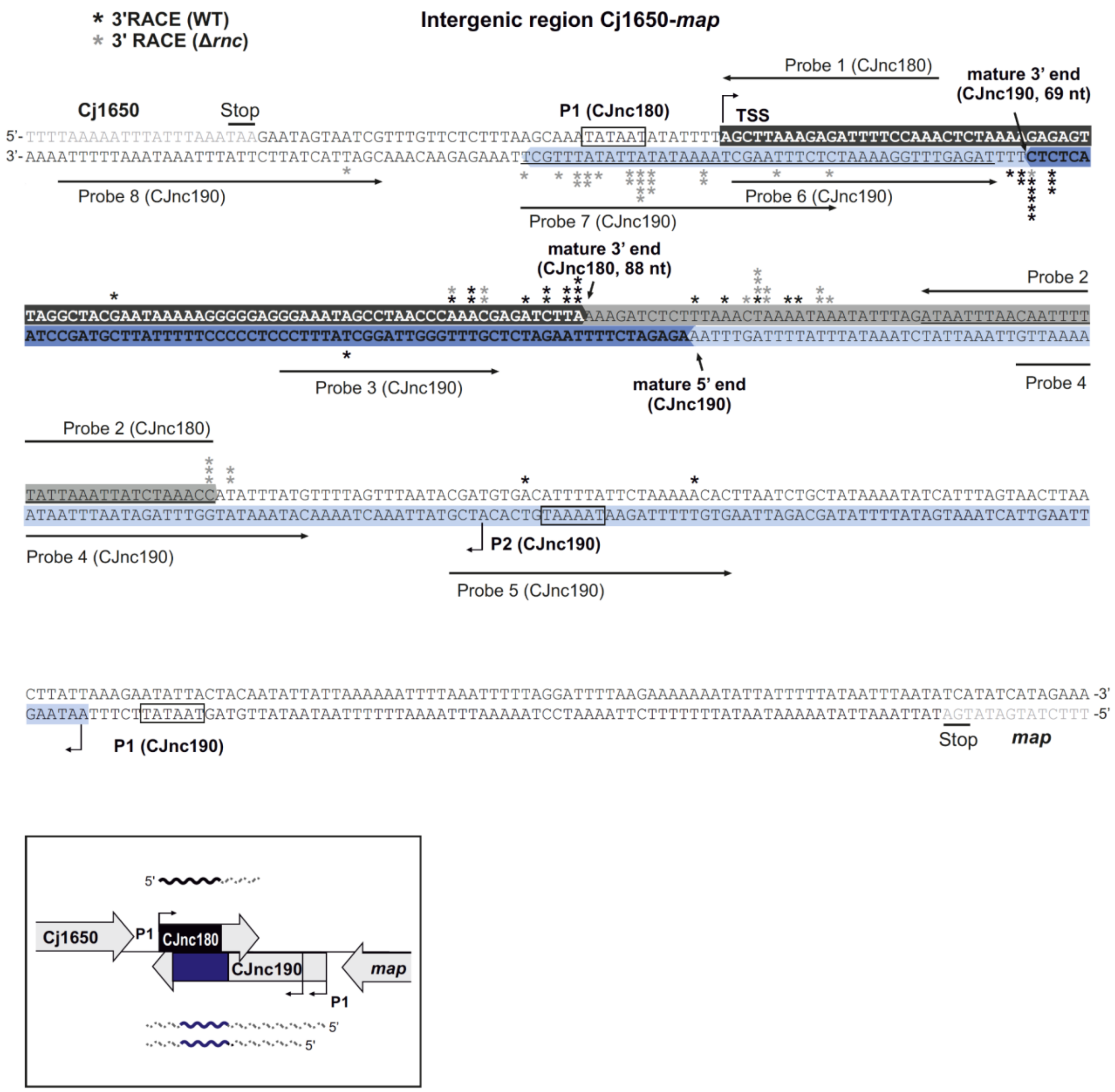


**Supplementary Figure S3. Identification of CJnc180/CJnc190 3' ends by RACE.** An RNA linker was ligated to 3′ ends of total RNA from WT or Δ*rnc*, followed by reverse transcription and PCR with linker/sRNA-specific primers. Cloned amplicons were sequenced to locate 3' end-linker junctions. Asterisks indicate 3′ ends detected in single RACE clones in WT (black) or Δ*rnc* (grey). Bent arrows: transcription start sites. Northern blot probe binding sites are underlined (Probes 1-8, see also **Supplementary Figure S7**). Dark grey/blue highlighted residues: mature CJnc180/190. Light grey/blue highlighted residues: mature CJnc180/190.

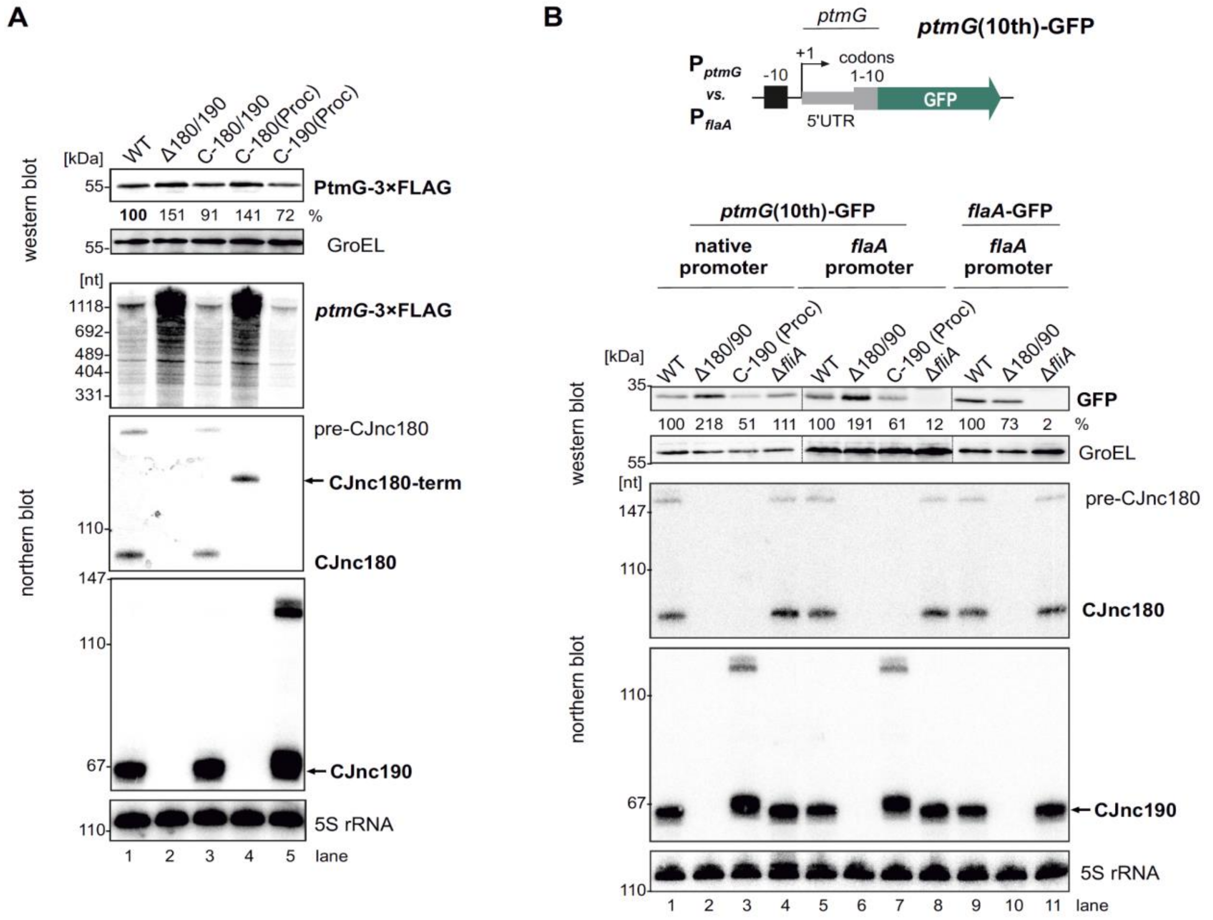


**Supplementary Figure S4. Post-transcriptional regulation of *ptmG* by CJnc190. (A)** CJnc190 is sufficient to repress *ptmG*. Related to main **Figure 1D**. Total protein and RNA samples were harvested from the indicated strains, and PtmG-3×FLAG levels were measured by western blotting with an anti-FLAG antibody, with GroEL as a loading control. Northern blot analysis confirmed the expression of a mature CJnc180-*rrnB* terminator fusion and a mature CJnc190-promoter P1 fusion (**Figure 1D**). "Mature" CJnc190 expressed in this context appears slightly longer than in WT. This is likely due to 3′ end differences, as its 5' end was confirmed to be the same as in WT by primer extension (**Supplementary Figure S2B**, lane 8). A putative transcriptional readthrough product (>110 nt) was also detected. **(B)** Promoter-independence of CJnc190 *ptmG* regulation. A *ptmG(10th)*-GFP translational reporter expressed from either the native *ptmG* promoter or the unrelated FliA ($\sigma^{28}$)-dependent *flaA* promoter was measured by western blotting in WT, Δ180/190, or C-190(Proc) strains. The 5′UTR (24 nt) and first ten codons of *ptmG* were fused to the second codon of GFP and introduced into the unrelated Cj0046 locus (Kim et al., 2008) in a Δ*ptmG* background. As a control, a $P_{flaA}$-*flaA* 5′UTR reporter was also included (lanes 9-11). CJnc190(Proc) was expressed from *rdxA*. The gene encoding $\sigma^{28}$ (*fliA*) was also deleted as a control (lanes 4, 8, 11). The PtmG(10th)-GFP reporter was detected with an anti-GFP antibody, and GroEL as a loading control. For all northern blots, probes for mature CJnc180/190 (CSO-0189/0185) and the 5′ end of the *ptmG* ORF (CSO-1666) or GFP (CSO-0789) were used, while 5S rRNA (CSO-0192) served as a loading control.

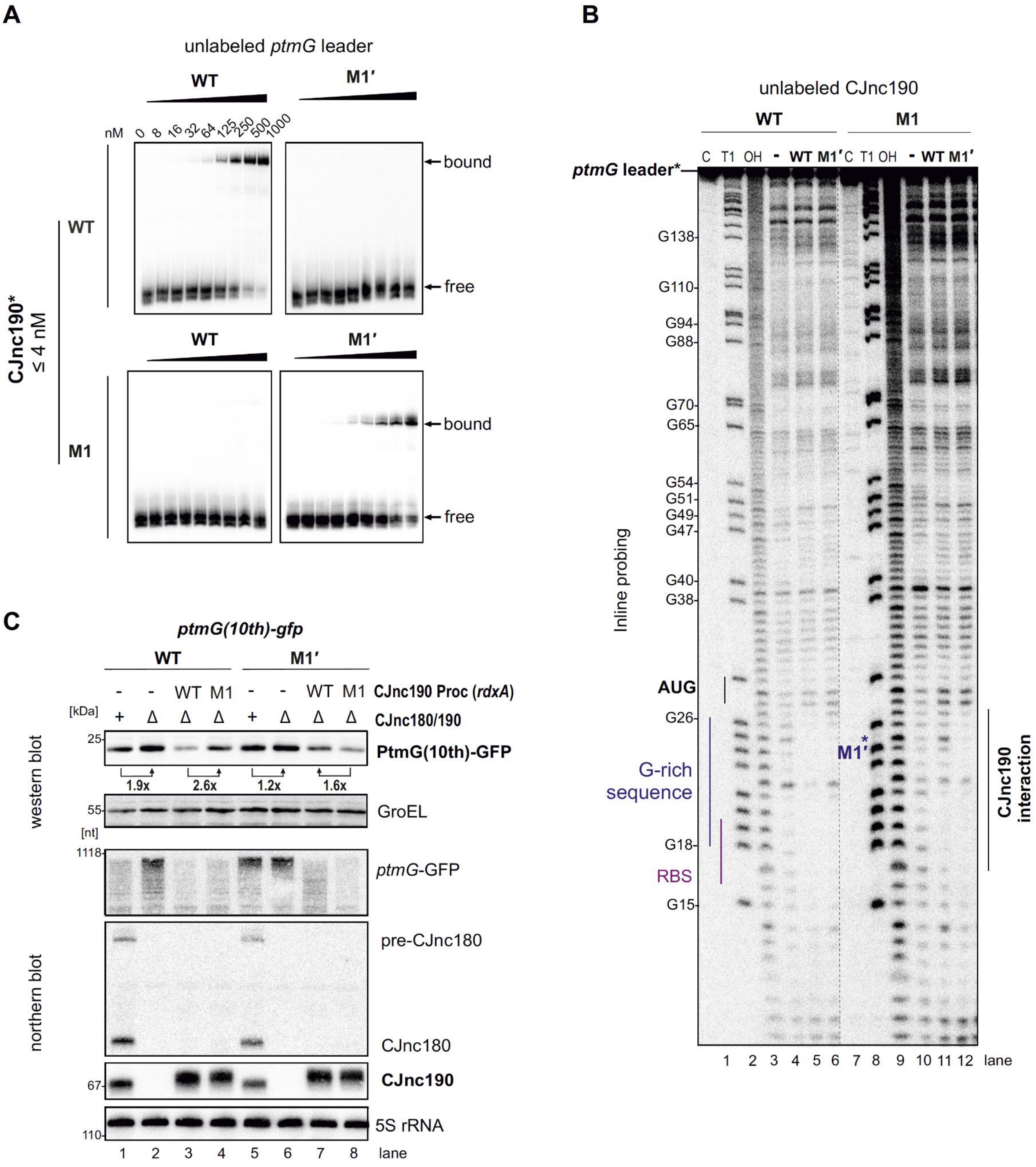


**Supplementary Figure S5. Direct repression of *ptmG* by CJnc190 via base pairing. (A)** *In vitro* gel mobility shift assay of mature CJnc190 binding to the *ptmG* 5′ UTR. $^{32}$P-Labeled (5′ end) CJnc190 *in vitro* transcript (WT/M1, see main Fig 2A) was incubated with increasing amounts of unlabelled *ptmG* leader (WT/M1′). Bound/free complexes were separated on native gels. **(B)** Inline probing of CJnc190 interaction with *ptmG*. $^{32}$P-labeled (5′ end) *ptmG* leader *in vitro* transcript (0.2 pmol) (WT or M1′, see main **Figure 2A**) was incubated with unlabeled CJnc190 (WT/M1) under mild alkaline conditions. Cleavage products were analyzed on denaturing gels. Lanes 1-3: C, no addition; T1, G residues (RNase T1); OH, all positions (alkaline hydrolysis). G residues are numbered with respect to the *ptmG* TSS (Dugar et al., 2013). The *ptmG* ribosome binding site (RBS), start codon (AUG), G-rich sequence, M1′ mutation, and CJnc190 interaction site are labeled. **(C)** Compensatory exchange analysis of *ptmG*:CJnc190 interaction. Related to main **Figure 2E**. A *ptmG(10th)-GFP* translational reporter (**Supplementary Figure S2B**) expressed from the Cj0046 locus was measured by western and northern blotting in a CJnc180/190 WT, Δ180/190, or C-190(Proc) (WT/M1, at *rdxA*) complemented background. The PtmG(10th)-GFP reporter was detected with an anti-GFP antibody. For northern blots, probes for mature CJnc180/190 (CSO-0189/0185) and the 5′ end of the *ptmG* ORF (CSO-1666) were used. GroEL served as a loading control for western blot analysis, while 5S rRNA served as a loading control on the northern blot (CSO-0192).

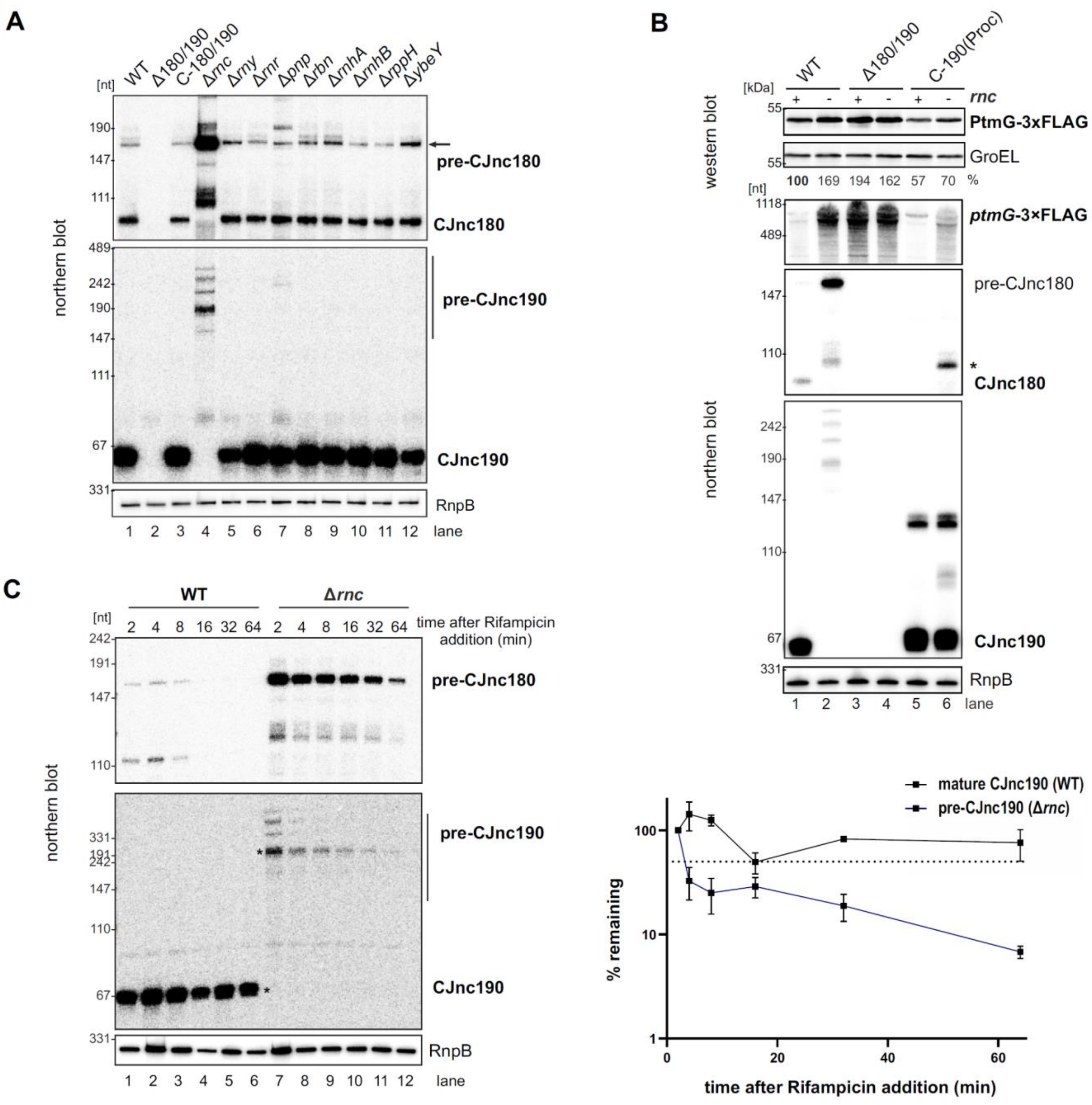


**Supplementary Figure S6. RNase III affects CJnc190 processing, stability, and *ptmG* regulation.** **(A)** Effect of deletions in genes encoding RNases and RNA-metabolism enzymes on CJnc180 and CJnc190 processing. Total RNA from WT and the indicated *C. jejuni* NCTC11168 mutant strains grown to mid-log phase was analyzed by northern blotting. Gene products are as follows: *rnc* - RNase III (Cj1635c); *rny* - RNase Y (Cj1209), *rnr* - RNase R (Cj0631c); *pnp* - PNPase (Cj1253); *rbn* - RNase BN (Cj1212c); *rnhA* - RNase HI (Cj1636c); *rnhB* - RNase HII (Cj0010c); *rppH* - RNA 5' pyrophosphohydrolase (Cj0581); *ybeY* - endoribonuclease YbeY (Cj0121). **(B)** RNase III affects PtmG expression via CJnc190 processing. Related to main Fig 3B. PtmG-3×FLAG protein and transcript were measured in WT, Δ180/190, and C-190(Proc) strains (Δ*rnc*/*rnc*+) by western and northern blot (CSO-1666), respectively. GroEL and RnpB (CSO-0497) served as loading controls for western and northern blots, respectively. The asterisk indicates a CJnc180 species still expressed from the C-190(Proc) construct. **(C)** RNA stability of CJnc180/190 in WT and Δ*rnc* measured after transcription inhibition by rifampicin during logarithmic growth. Probes specific for processed CJnc180/190 (CSO-0189/0185) or the 5′ end of the *ptmG* ORF (CSO-1666) were used for all northern blots, while RNase P RNA (RnpB, RNase III-independent) was probed as a loading control (CSO-0497). (*Right*) Quantifications of bands with asterisks (*left*) with standard errors calculated from two independent replicates. Dashed line indicates 50% remaining.

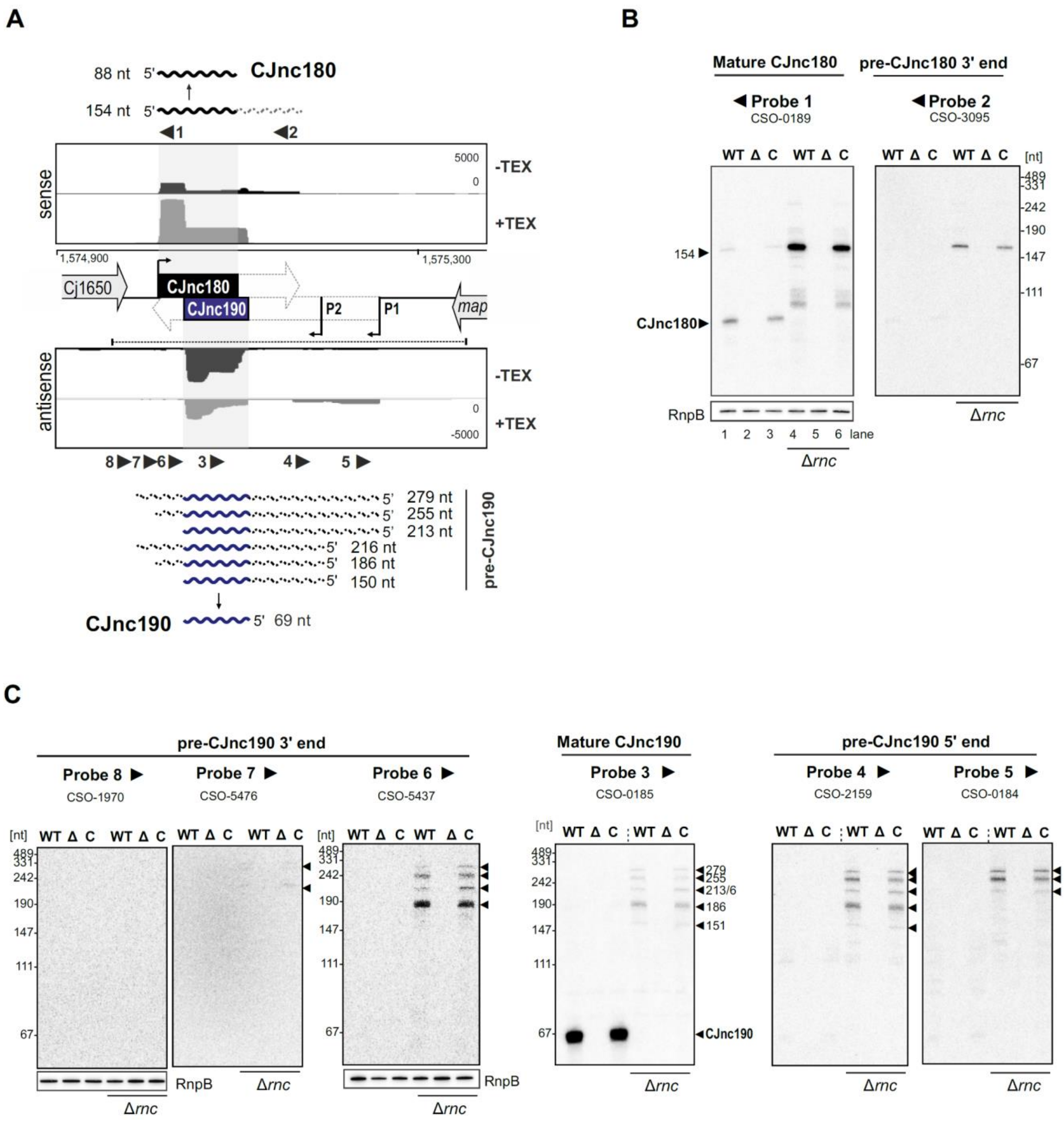


**Supplementary Figure S7. Northern blot analysis of mature and precursor CJnc180 and CJnc190. (A)** Approximate location of northern blot probes used in panels B and C and schematic of the detected CJnc180 and CJnc190 species in WT and Δ*rnc* backgrounds. See also **Supplementary Figure S3** for probe binding sites. **(B & C)** Northern blot analysis of CJnc180 and CJnc190 for CJnc180/190 WT, deletion, and complementation strains in total RNA. Analysis was performed in both *rnc*+ and Δ*rnc* backgrounds. All panels are the same RNA samples. Probings with CSO-0189, CSO-3095 (panel B) and CSO-0185, CSO-2159, and CSO-0184 (panel C) are on the same blot. CSO-5437 and CSO-5476/CSO-1970 were used to probe two additional blots with the same RNA samples. RnpB was also probed as a loading control (CSO-0497), and is shown only once for each blot.

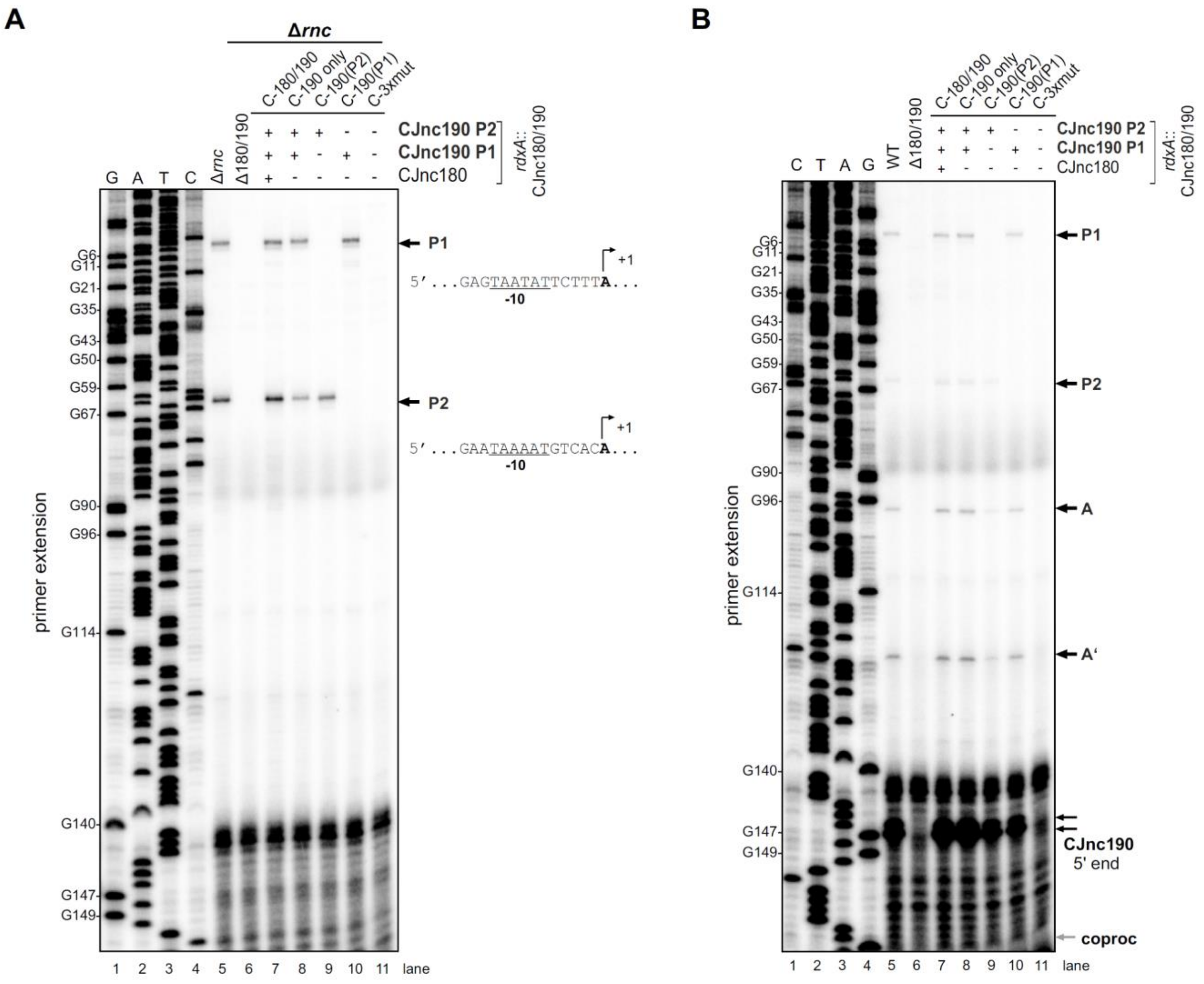


**Supplementary Figure S8. Mapping of CJnc190 mature and precursor 5’ ends in promoter mutants. (A)** Primer extension-based detection of pre-CJnc190 5’ ends for WT and promoter mutant strains in a Δ*rnc* background. Five-prime ends related to promoters P1 and P2 are indicated (*right*) together with their putative -10 boxes. Related to main **Figure 3C**. C-3×mut: strain complemented with CJnc180/190 region carrying point mutations in all three predicted promoter -10 boxes. **(B)** Primer extension analysis of CJnc190 5’ ends in an RNase III+ background. Putative precursor 5’ ends (A/A’, **Figure 5A**) and where a putative co-processing site (coproc, see **Figure 6A**) would be visible are indicated on the right. For both panels, total RNA was subjected to primer extension with a probe annealing to mature CJnc190 (CSO-0185). Samples for both panels were run on the same gel with the same sequencing ladder generated from the same probe and the CJnc180/190 region of *C. jejuni* NCTC11168 (lanes 1-4), but the gel was cut and separated into Δ*rnc* and WT background sample panels for clarity. G residues (numbered based on the P1 primary transcript) are indicated on the left.

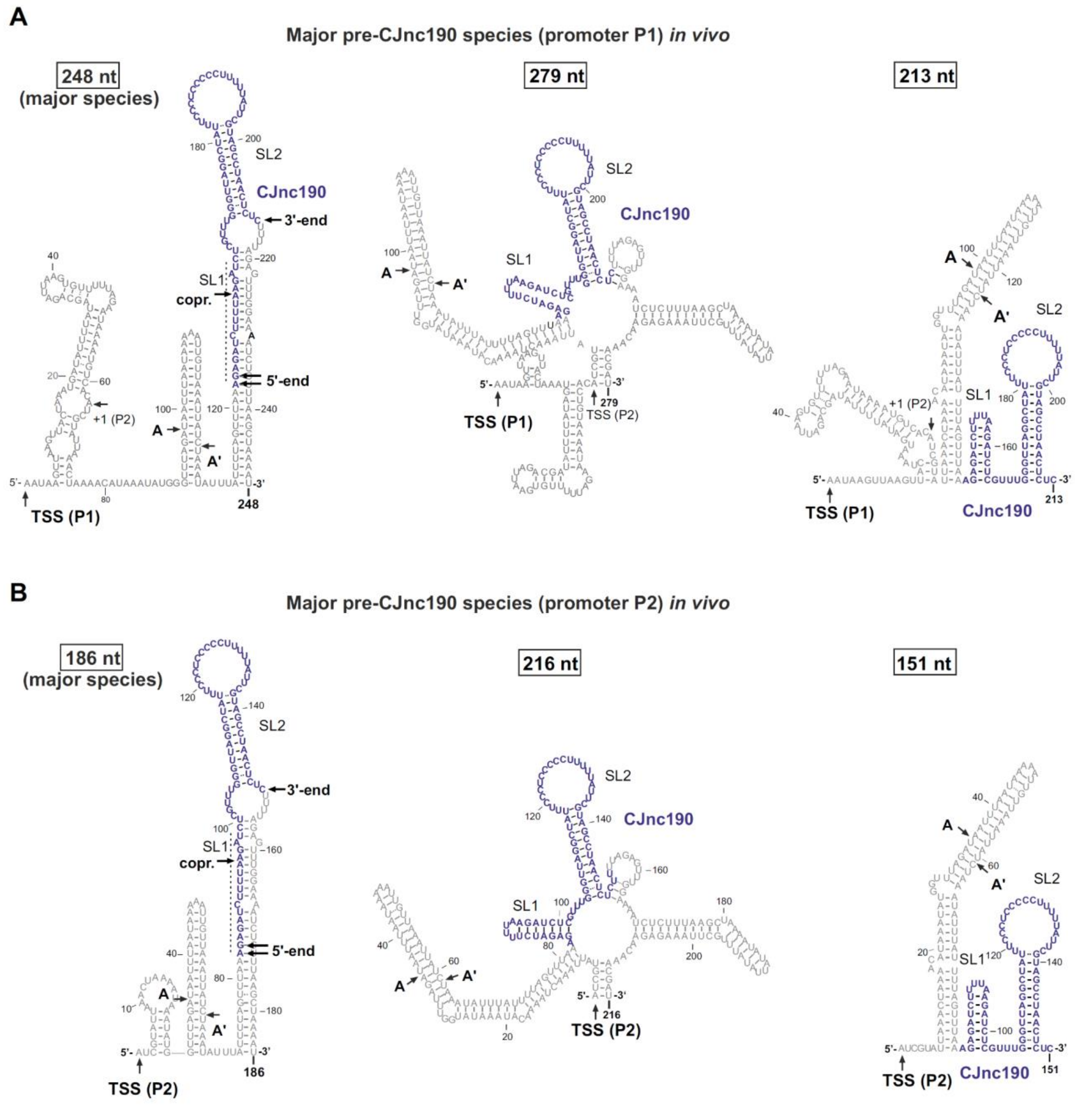


**Supplementary Figure S9. Predicted secondary structures of diverse pre-CJnc190 species *in vivo*.** Secondary structures of CJnc190 species transcribed from promoter P1 (**A**) and P2 (**B**) as suggested by primer extension and 3′RACE (**Supplementary Figure S2 & S3**), as well as northern blotting (**Figure 3A**), in Δ*rnc*. The most abundant species for P1 and P2 (based on analysis in the presence of CJnc180) are indicated. Secondary structures were predicted using the RNAfold server of Vienna RNA tools (Lorenz et al., 2011). (A/A′): potential intermediate RNase III cleavage sites (see **Supplementary Figure S10**). (copr.): Potential co-processing site with CJnc180 (**Figure 6A**).

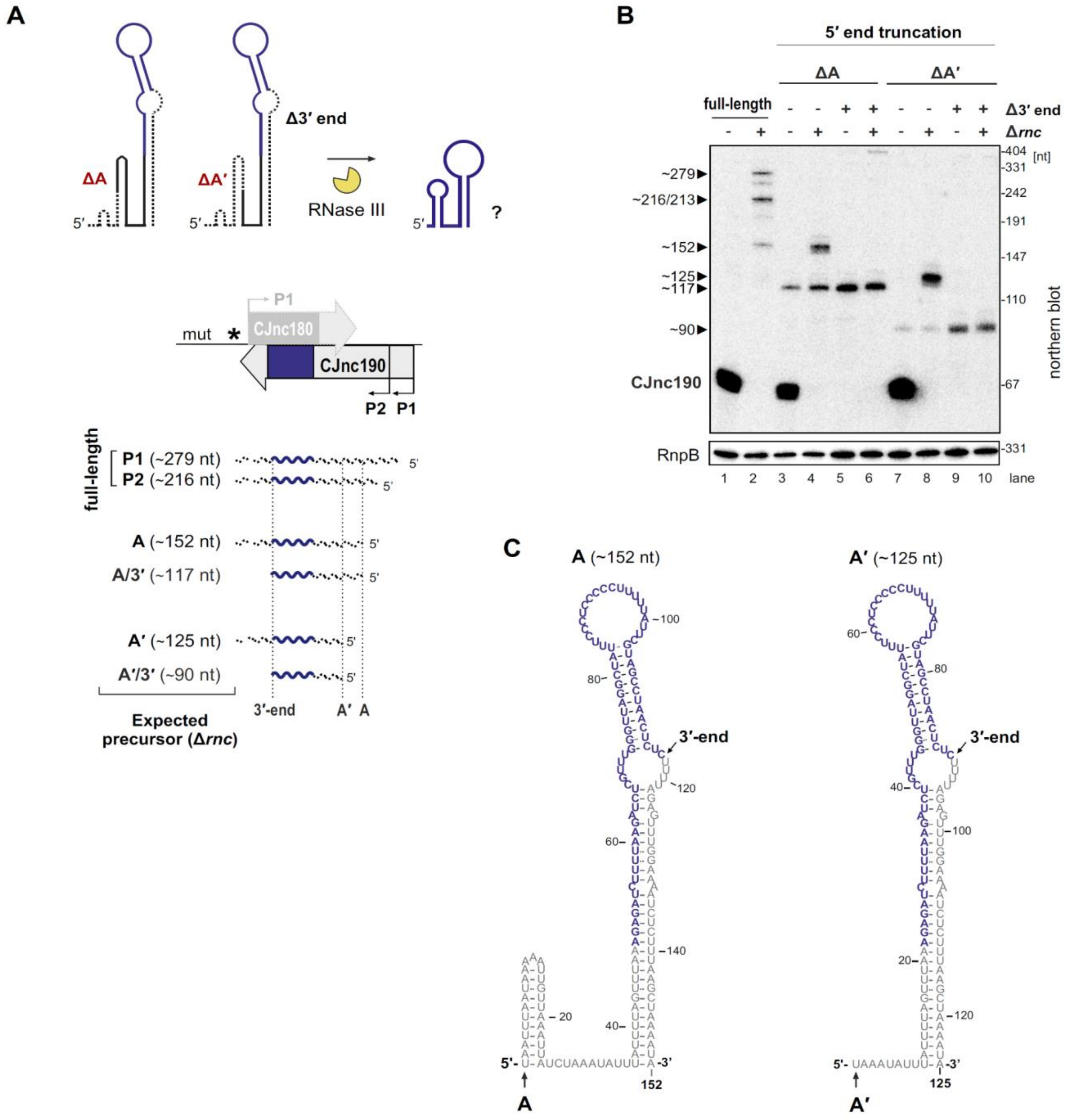


**Supplementary Figure S10. Processing of pre-CJnc190 truncated in a 5'-end hairpin. (A)** Strategy for testing requirement of the 5' end hairpin for processing. CJnc190 versions (5'-truncated at positions A/A' (ΔA/ΔA') or 3'-truncated at mature sRNA end (Δ3')) were expressed from *rdxA* of Δ180/190 (without CJnc180). Expected precursor lengths are shown. **(B)** Northern blot analysis of CJnc190 processing in strains expressing truncated CJnc190 versions outlined in A. CJnc190 expression and processing was measured by northern blotting with a probe binding the mature CJnc180/190 sRNAs (CSO-0189/0185). RnpB (CSO-0497) was probed as a loading control. **(C)** Predicted secondary structures of pre-CJnc190 (promoter P2, based on 186 nt precursor in **Figure 5A**) species truncated at 5′ positions A or A′. Structures were predicted using the RNAfold server of Vienna RNA tools (Lorenz et al., 2011).

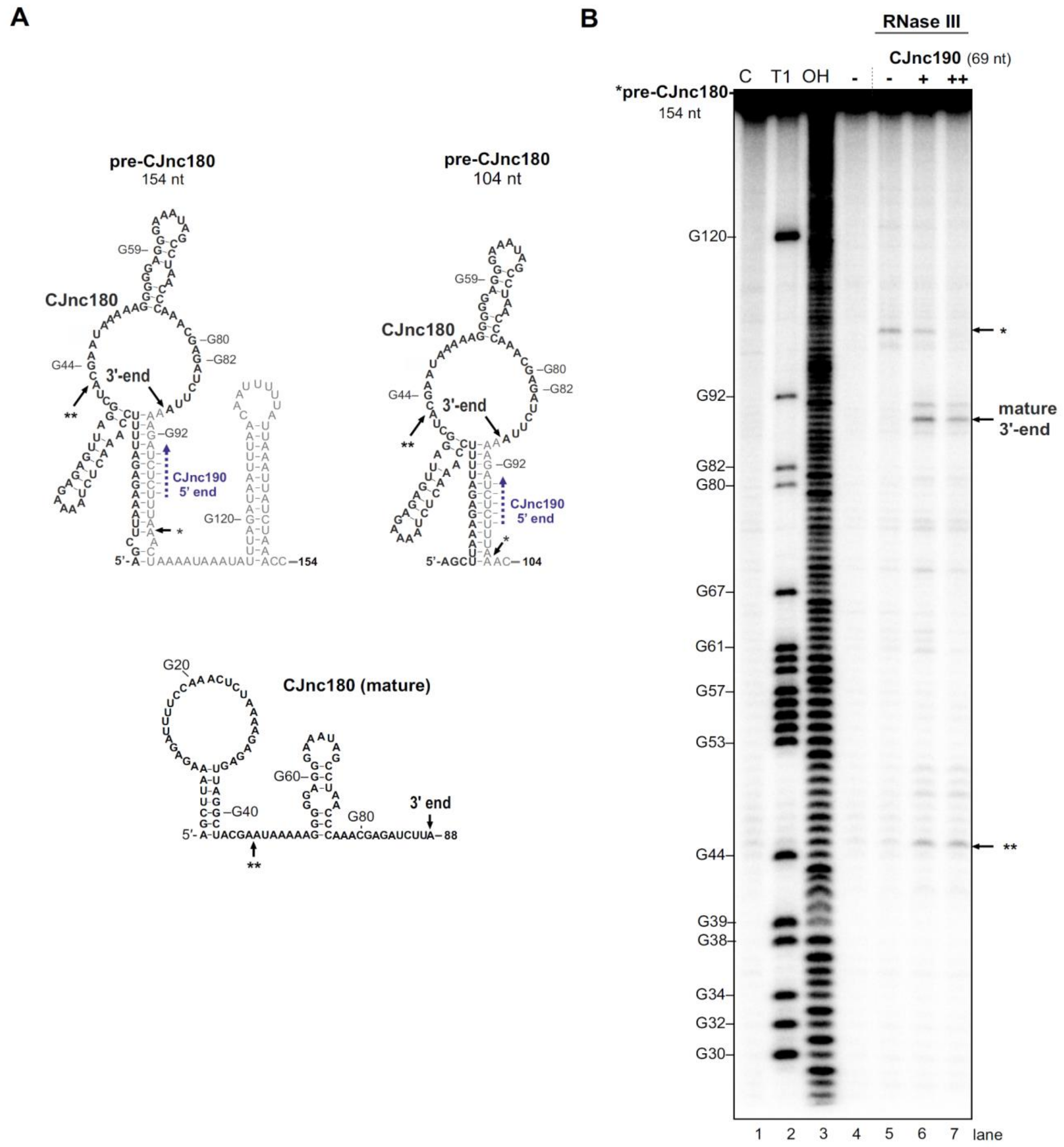


**Supplementary Figure S11.** ***In vitro*** **RNase III processing of pre-CJnc180. (A)** Predicted secondary structures of pre-CJnc180 versions (154 and 204 nt) and mature CJnc180 (RNAfold (Lorenz et al., 2011)). The *in vivo*-detected 3' end is indicated. *: CJnc190-independent *in vitro* RNase III cleavage site (panel B). **: CJnc190-dependent *in vitro* RNase III cleavage site. Blue dashed arrow: Region of potential base-pairing with 5′ end of mature CJnc190. **(B)** *In vitro* RNase III cleavage of pre-CJnc180 alone or in the presence of unlabeled mature CJnc190. A pre-CJnc180 *in vitro* transcript ($^{32}$P 5' end-labeled, 0.2 pmol) was treated with 1/625 U RNase III. Where indicated, unlabeled mature CJnc190 RNA was added (+: 0.2 pmol, ++: 2 pmol). Cleavage products were separated on denaturing gels. Lanes 1-3: C, no addition; T1, G residue ladder (RNase T1); OH, all positions (alkaline hydrolysis). Related to main **Figure 6B**.

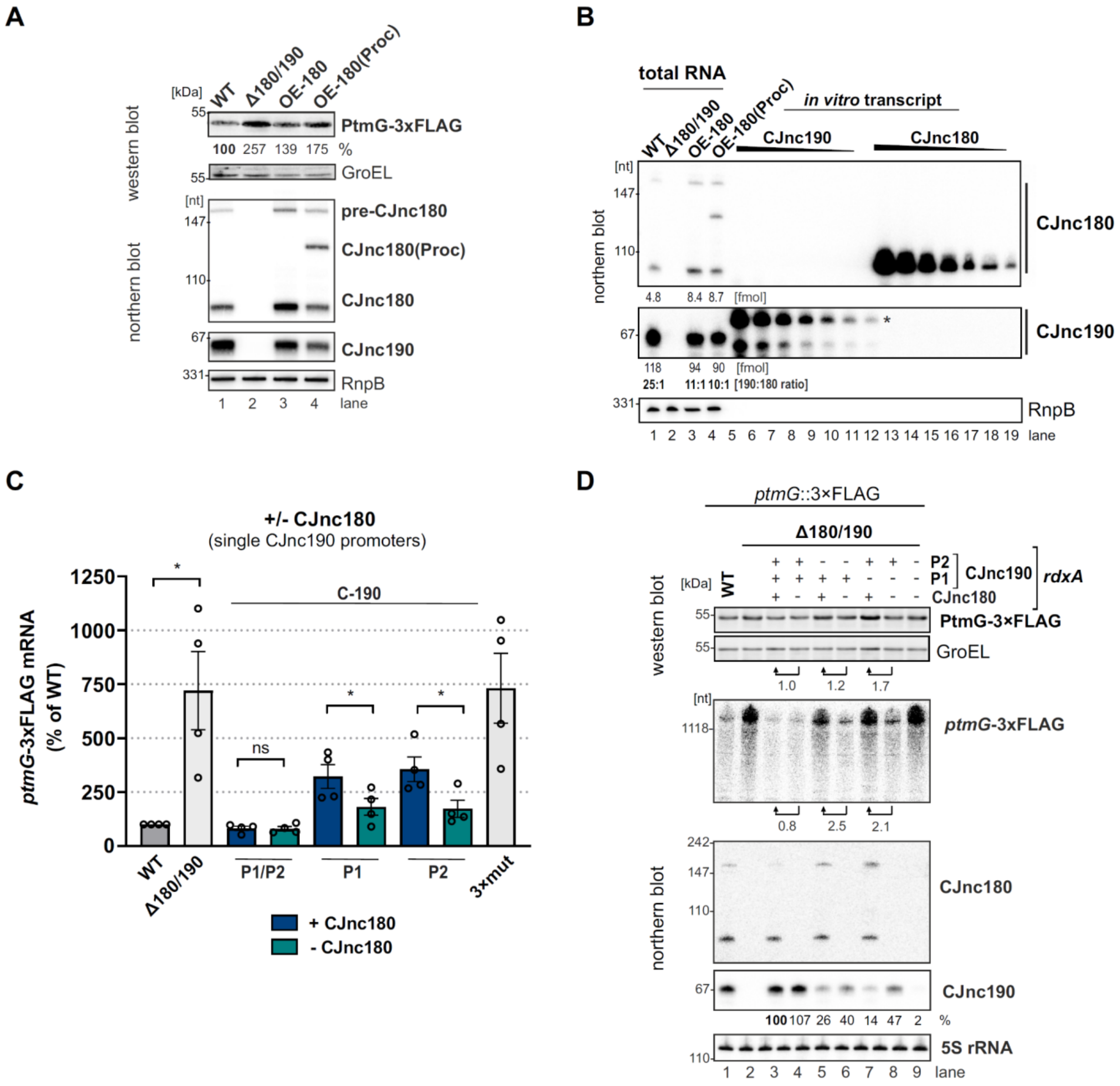


**Supplementary Figure S12. Modulation of *ptmG* regulation via CJnc180 expression and different CJnc190 promoters. (A)** Expression of a second copy of CJnc180 or CJnc180(Proc) derepresses *ptmG*. Levels of PtmG-3×FLAG protein were measured by western blot in the indicated strains in log phase. GroEL was detected as a loading control. Related to main **Figure 7A**. **(B)** Estimation of relative CJnc180 and CJnc190 levels in a WT or OE-CJnc180 background in log phase. CJnc180 and CJnc180 levels in total RNA were determined by comparison to serial dilutions of *in vitro* transcripts by northern blot analysis. The regions of the lanes that were quantified for CJnc180 and CJnc190 are indicated by vertical bars on the right. Related to main **Figure 7A**. **(C)** Absence of CJnc180 de-represses *ptmG* when CJnc190 is expressed from a single promoter. Levels of *ptmG*-3×FLAG protein were measured by northern blot in the indicated strains in log phase. 3×mut: Δ180/190 complemented with CJnc180/190 carrying point mutations in all three validated promoters. Error bars: SEM from four independent replicates. Student's unpaired *t*-test vs. WT: *: $p<0.05$, ns: not significant. Related to main **Figure 7B**. **(D)** Representative western/northern blot for the effect of the CJnc180 antagonist absence or presence on PtmG levels in log phase. Related to main **Figure 7B and panel C**. For northern blots, probes for mature CJnc180/190 sRNAs (CSO-0189/0185) and the 5′ end of the *ptmG* ORF (CSO-1666) were used. GroEL and 5S rRNA (CSO-0192) were detected as loading controls for protein and RNA samples, respectively. PtmG-3×FLAG fold-changes were calculated with GroEL-normalized measurements.

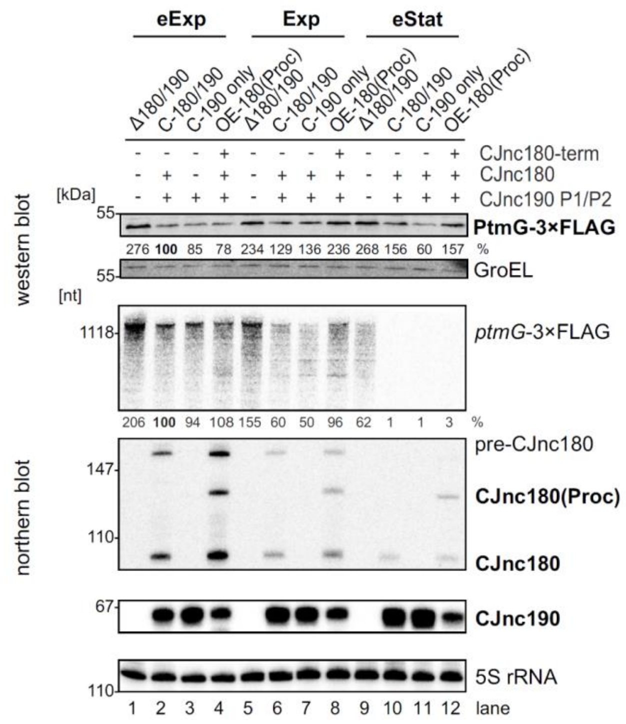


**Supplementary Figure S13. Modulation of *ptmG* regulation via CJnc180 expression at different growth phases.** Related to main **Figure 7D**. Lag: lag phase, eExp: early exponential, Exp: exponential, eStat: early stationary phase ($OD_{600}$ 0.1, 0.25, 0.5, and 0.9, respectively). Northern blot probes for mature CJnc180/190 sRNAs (CSO-0189/0185) and the 5' end of the *ptmG* ORF (CSO-1666) were used, as well as CSO-0192 (5S rRNA) as a loading control. +/- denotes whether the indicated sRNA is expressed in the strain.

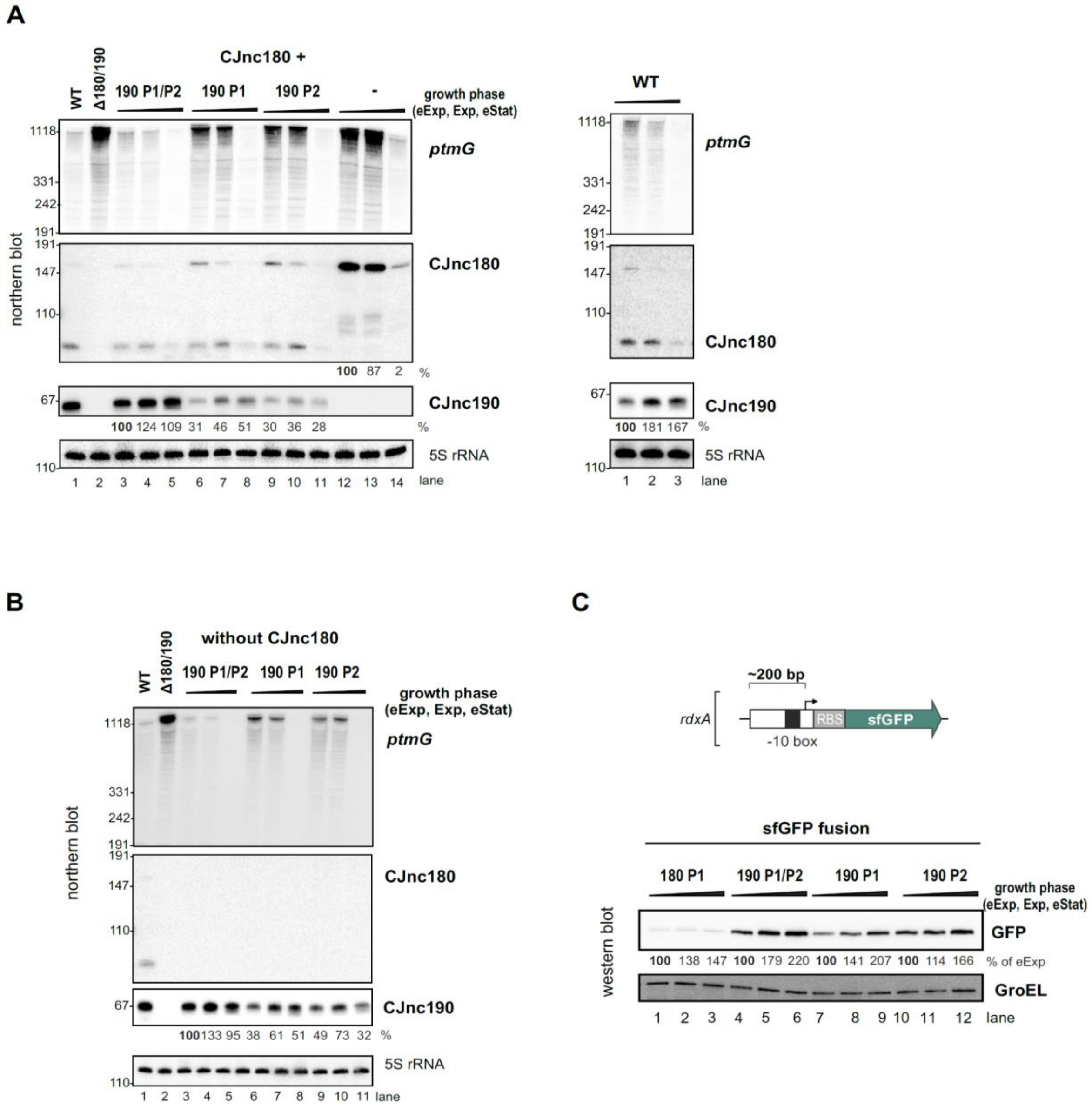


**Supplementary Figure S14. CJnc180 and CJnc190 levels and promoter activity at different growth phases. (A & B)** Levels of CJnc190 expressed from promoter P1 or P2, as well as *ptmG*, at different growth phases. Levels were determined in the presence (A) or absence (B) of CJnc180. Promoter point mutant strains (see main **Figure 3C**) were used. **(C)** Expression of sfGFP transcriptional reporters for CJnc180 P1 and CJnc190 P1/P2, P1, or P2 at different growth phases. Approximately 200 bp upstream and 10 bp downstream of the CJnc180, CJnc190 P1, and CJnc190 P2 TSSs were fused to sfGFP with an RBS from the unrelated *hupB* gene to generate CJnc180 P1, CJnc190 P1, and CJnc190 P1/P2 strains. Site-directed mutagenesis (see **Supplementary File 1 - Table S5** for details) was used to inactivate CJnc190 P1 in the P1/P2 strain to generate a CJnc190 P2-only reporter. GFP levels in cells harvested from the indicated growth phases were detected using an anti-GFP antibody, with GroEL as a loading control. Black triangles above the all blots indicate increasing growth phase (eExp, Exp, Stat). Northern blot probes mature CJnc180/190 (CSO-0189/0185) and the 5’ end of the *ptmG* ORF (CSO-1666) were used, as well as 5S rRNA (CSO-0192) as a loading control.
